# Extensional Flow Microfluidics Reveal Size-Dependent Core-Shell Mechanical Heterogeneity in Pancreatic Tumor Spheroids

**DOI:** 10.64898/2026.11.04.764045

**Authors:** Sara Ghanbarpour Mamaghani, Timothy M Costa, Ethan C Wagner, Soroush Kargar, Jonathan P Celli, Joanna B Dahl

## Abstract

Understanding the mechanical behavior of cancer cell aggregates is important for investigating metastatic potential and developing effective therapeutics. Tumor cell clusters are major contributors to metastasis and can undergo extreme deformation to migrate through capillaries smaller than the cluster size, suggesting complex multicellular mechanics. Despite their clinical significance, the mechanics of multicellular spheroids remain underexplored. In this study, the mechanical behavior of spheroids formed from the pancreatic cancer cell line PANC-1 was investigated using a cross-slot microfluidic device. Suspended spheroids from two size ranges were stretched in non-contact extensional flow (small spheroids mostly 40 < *D*_0_ < 75 µm, large spheroids mostly 90 < *D*_0_ < 145 µm). Digital image correlation analyses of intra-spheroid strains revealed strong deformation differences for small and large spheroids while analyses based on 2D projected contours of the spheroid shape did not show trends. Smaller spheroids deformed relatively homogeneously and exhibited significant overall stretching. In contrast, larger spheroids displayed heterogeneous deformation characterized by a stiff central core and a deformable outer shell. To investigate the origin of these mechanical differences, cellular morphology and the expression of stiffness-associated proteins, including vimentin and E-cadherin, were assessed from confocal microscopy. Larger spheroids display significantly lower nuclear sphericity than small spheroids, indicating more cellular packing with increasing spheroid size. Additionally, vimentin expression was concentrated in the outer shell of large spheroids, whereas small spheroids exhibit vimentin expression throughout the aggregate. Thus individual cell morphology and cytoskeletal networks seemed to contribute to the core-shell biomechanical behavior of the large spheroids and lack thereof for small spheroids. Taken together, this study suggests that as spheroids grow, they transition from an irregular and disorganized cluster at low cell numbers to a coordinated and hierarchical structure at sufficient cell numbers that corresponds to diameters ≳ 100 µm.

## 1 Introduction

Understanding the mechanical behavior of cell aggregates is important in a range of biological and biomedical contexts, including tissue engineering [1], tissue repair [2], cancer metastasis [3], and therapeutic applications [4, 5]. In tissue engineering, biomaterials containing cells and extracellular matrix (ECM) are used for wound healing and tissue regeneration, where mechanical properties of tissues play a critical role in clinical applicability [6]. In this context, engineered cell aggregates like tumors serve as simplified model systems for tissues, enabling the study of cell-cell interaction and 3D effects on mechanics. Cell aggregates are also central to cancer progression and metastasis. Circulating tumor cell clusters have been shown to have a significantly higher metastatic potential than single cells [7, 8]. These clusters can deform through capillaries smaller than their own diameter while still maintaining viability [3], and an improved understanding of their mechanics could facilitate better cancer treatments. In clinical settings, mechanical properties of single cells have provided complementary biophysical information to biochemical markers for more robust cellular phenotyping [9] and label-free sample enrichment [10], leading to FDA-cleared clinical diagnostic devices that process patient blood samples (e.g., the Cytovale In-tellisep Test for early sepsis detection [11] and the CelLBxHealth’s Parsortix Platform [12] that enriches for circulating tumor cells and clusters). Similarly, better understanding of cellular aggregate mechanics could improve spheroid and organoid biomedical clinical translation. However, compared with single cells, the mechanical properties of cell aggregates remain much less studied.

The various methods used to characterize the mechanical properties of cellular spheroids vary in where the deforming force is applied and on what timescale [1]. Traditional micromechanical techniques like atomic force microscopy (AFM) [13], micropipette aspiration (MPA) [14], and parallel plate compression [15, 16] have been used to test spheroid mechanics from seconds to hours timescales. These studies have revealed that spheroids are viscoelastic materials: cells respond to force by changing their own stiffness [13], and they rearrange under force and do not fully bounce back [15]. For AFM and MPA in which the deforming force is applied on a small portion of the spheroid surface, it is possible that the mechanical behavior of the local area around the applied force dominates the measurement, making it challenging to capture the bulk spheroid mechanical behavior or mechanical heterogeneities around the spheroid.

Recently, microfluidic approaches have assessed spheroid mechanics at higher throughputs and faster timescales. Boot et al. [17] developed a microfluidic platform adapted from micropipette aspiration technique to measure the seconds-scale viscoelastic properties of tens of spheroids per day in parallel. Panhwar et al. [18] used microfluidic virtual straight channels to deform spheroids with shear stresses on millisecond timescales and estimated the spheroids’ bulk Young’s moduli, which were interestingly 10 times softer than single cells. Pandey et al. [19] used a microfluidic compression device to measure the viscoelastic properties of tumor spheroids, finding that relaxation dynamics could distinguish malignant from non-tumorigenic spheroids. These studies introduced their microfluidic techniques using proof-of-concept measurements on spheroids from a relatively narrow size range with diameters larger than 100 µm (120–250 µm [17], approximately 130 µm [18], and 143–154 µm [19]). Given that spheroid mechanics is governed by multiple factors including the stiffness of individual cells and the strength of cell-cell junctions [20], and that cell volume, cell stiffness, and cell-cell junctions can vary spatially within the spheroid volume [21], one can expect that spheroid mechanics will change with spheroid size as cell and cell-cell junction numbers change and the multicellular system dynamically adapts. Furthermore, *>*100 µm spheroid size ranges are also larger than the sizes of circulating tumor cell clusters isolated from patient samples, which contain 2–20 cells and estimated to be on the order of tens of micrometers in diameter [3]. Little is known about the continuum of biomechanical behavior from single cells to tissue-like spheroids and the implications for hierarchical cellular tissue organization, wound healing, diseases, and potential clinical applications.

To investigate the effect of spheroid size on their mechanics, we examined the deformation behaviors of PANC-1 pancreatic cancer cell spheroids stretching in non-contact microfluidic extensional flow for two different size ranges (average diameters 55 and 135 µm). For comparison, the same deformation experiments were performed on PANC-1 single cells to assess differences in mechanical response between individual cells and multicellular aggregates. Analysis of cell-scale strains at the spheroid near-surface revealed that larger spheroids above a critical size threshold of approximately 90 µm had a heterogeneous core-shell deformation pattern, with a softer outer cell layer deforming more than the spheroid core. In contrast, smaller spheroids deformed a similar amount throughout the spheroid. Single cells deformed less under similar extensional hydrodynamic forces compared to both small and large spheroids made up of that same cell type. To probe the physical and biological factors underlying the observed mechanics differences, we analyzed the cellular morphologies within spheroids and examined the expression and distribution of two cytoskeletal and adhesion-related proteins, vimentin and E-cadherin, associated with tissue integrity and mechanical stiffness. Overall, our results suggest that high-throughput microfluidic extensional mechanical testing of multicellular spheroids can pull out mechanical inhomo-geneities and efficiently explore the variation in mechanical behavior across these complex microscale tissue mimics.

## 2 Results and Discussion

### 2.1 Microfluidic Extensional Flow Technique

#### 2.1.1 Spheroid Characterization

PANC-1 spheroids were cultured for four days prior to microfluidic experiments to allow for formation of cohesive cell aggregates while limiting the possibility of a necrotic core that could affect spheroid mechanics (see Experimental section for more details). Two spheroid size groups were fabricated by varying the initial cell seeding density: small spheroids with a target average diameter of 50 and large spheroids targeting 100 µm average diameter. To assess the timing of the onset of necrotic core formation, the large spheroids were monitored for growth and viability over 15 days (**Figure 1**). Spheroid effective diameter as measured from the 2D projected spheroid area followed a monotonically increasing trend, as expected (**Figure 1A**). After one day of culture, the cells aggregated to form small clusters. By day 4, the spheroids reached an average diameter around 140 µm, greater than the target size of 100 µm diameter. This discrepancy between the target and actual average large spheroid size may be due to mild settling and flattening of the spheroids under gravity, which increases their apparent diameter in 2D projections from the spherical shape assumed when calculating the initial cell seeding density. The necrotic core area measured from ethidium bromide staining was negligible for spheroids cultured up to five days, while beyond a week of culturing the interior of the spheroids consisted of primarily dead cells (**Figure 1B**). (Representative live-dead stained spheroid images are found in Supplementary Figure S1.) Because the spheroids used in the microfluidic experiments were cultured for only 4 days, the very small necrotic area measured at this time point indicates that the mechanical measurements obtained from these spheroids primarily represent the behavior of viable spheroids consisting of mostly living cells.

**Figure 1:**
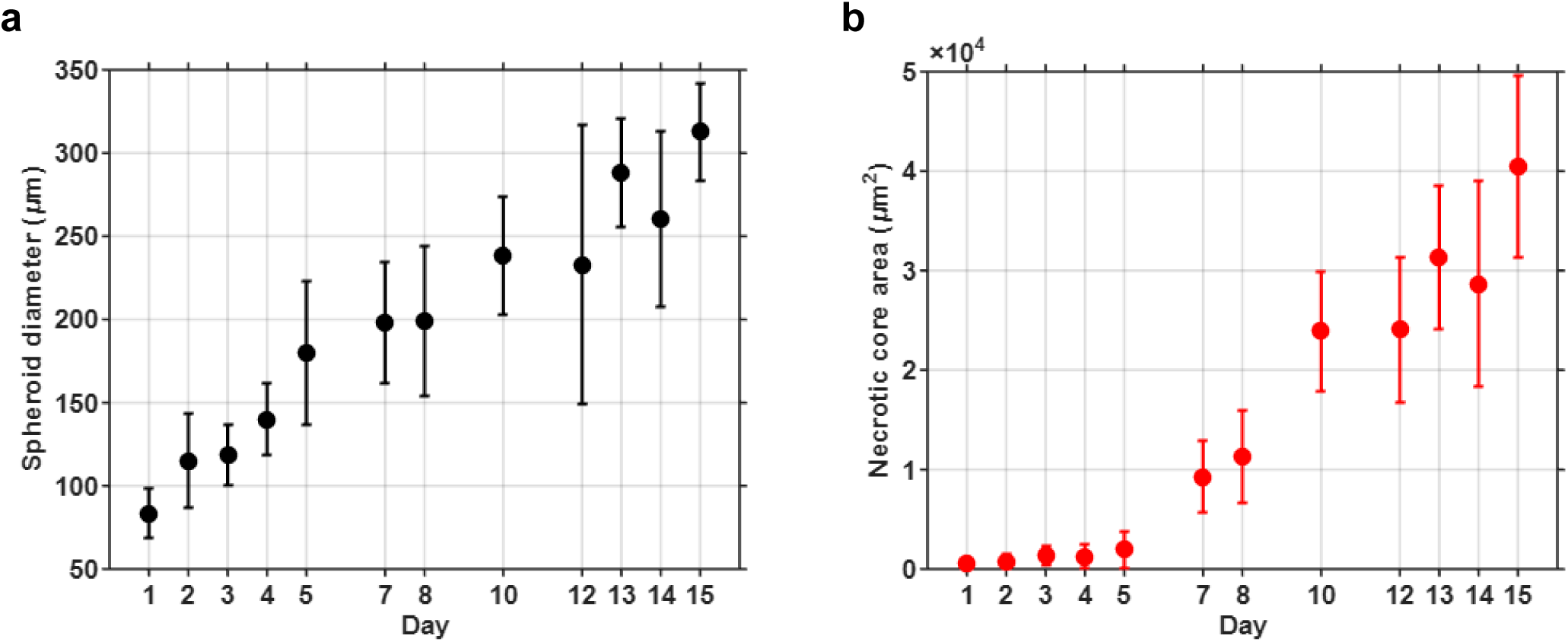
Spheroid size and viability characterizations. **(a)** Spheroid diameter and **(b)** spheroid necrotic core area over culture days 1–15. Dots represent the mean and error bars represent the standard deviation. Day 0 corresponds to the day of cell seeding. On each culture day, data from four independent experiments were combined for an average of 53 individual spheroids were measured for diameter and 60 individual spheroids were assessed for viability. Spheroid size increased monotonically with culture time. The necrotic core was of negligible size up to 5 days of culturing.

#### 2.1.2 Stretching of Spheroids and Cells in Microfluidic Extensional Flow

Microfluidic uniaxial deformation experiments were performed on individual PANC-1 spheroids using a cross-slot extensional flow microfluidic device with sheath flow focusing (**Figure 2**). To provide a basis for comparison between spheroids and their constituent cells, deformation experiments were also conducted on single PANC-1 cells. Extensional flow was generated in the central cross-slot region where the sample stream (cells or spheroids suspended in the viscous buffer solution) impinged on blank buffer stream entering the opposite inlet channel. The viscous buffer consisted of 0.8% w/v methylcellulose in DPBS (1×) containing calcium. **Figure 2A** depicts a schematic of the extensional flow microfluidic device and the parameters influencing the viscous fluid forces applied to the spheroid. The technique is illustrated through a representative single cell, small spheroid, and large spheroid results showing superimposed contours from the selected movie frames are shown (**Figure 2B**) along with the corresponding time histories of cell/spheroid speed and deformation (**Figure 2C**). Full time-lapse movies for these three experiment groups are provided in **Movies S1–S3** (Supporting Information).

**Figure 2:**
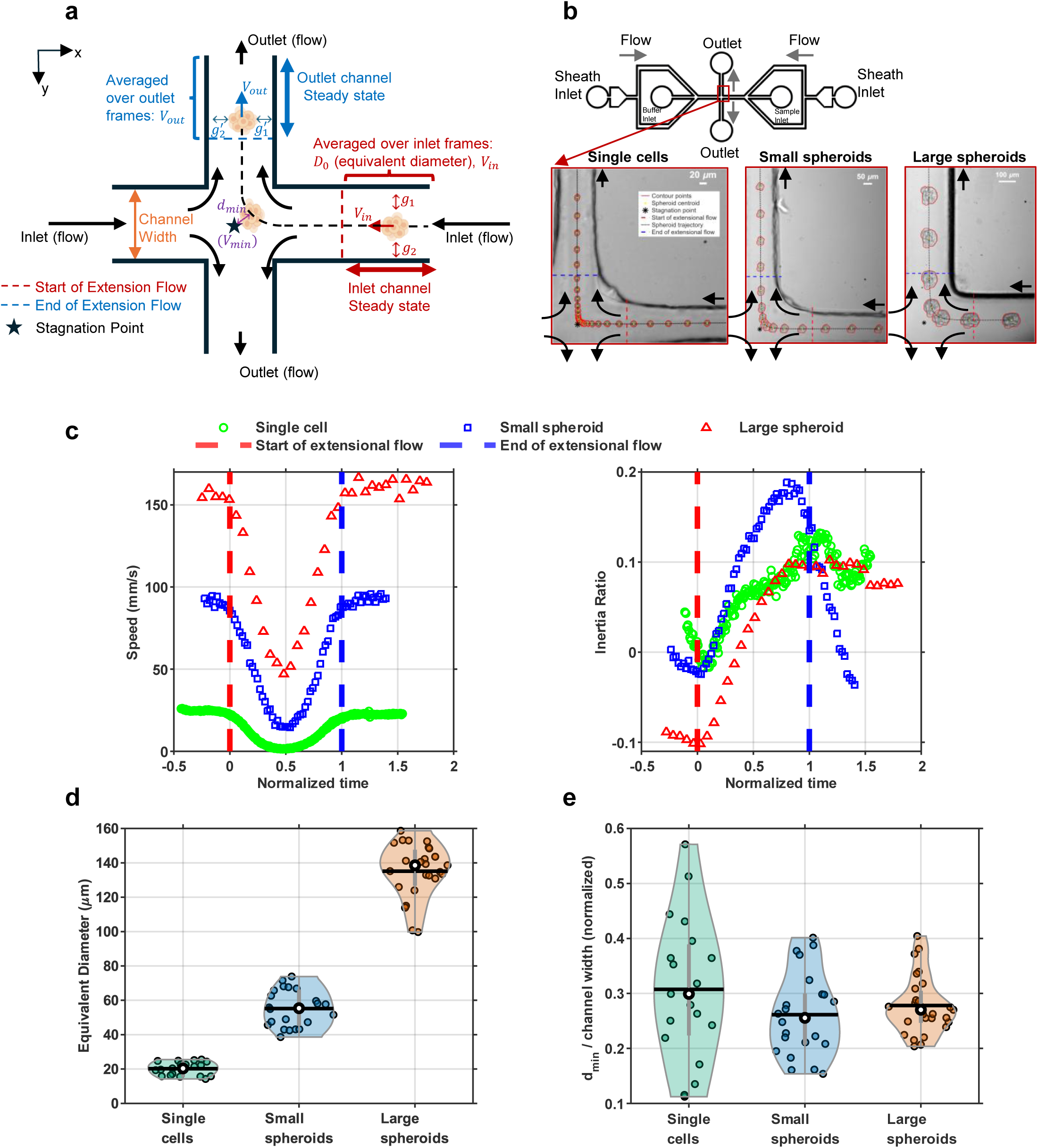
Microfluidic cross-slot technique to stretch spheroids of different sizes and single cells in similar extensional flow conditions. **(a)** A technique schematic showing a spheroid stretching under extensional flow. **(b)** Select movie frames for a representative single cell (*D*_0_ = 17.2 µm), small spheroid (47.5 µm), and large spheroid (99.7 µm) under approximately matched extensional hydrodynamic stress conditions (*µ*Ω = 71±11 Pa). **(c)** Time histories of the center-of-mass speed and 2D contour inertia ratio for the representative cell and small and large spheroids. Time is normalized by the duration each object spent in the extensional flow region (between the dashed red and blue lines): 49.77 ms for the single cell, 14.92 ms for the small spheroid, and 5.09 ms for the large spheroid. **(d)** Equivalent diameter *D*_0_ distributions for all single cells (mean 20.2 µm; range 14.2–25.5 µm; *N* = 19), small spheroids (mean 55.2 µm; range 38.6–73.9 µm; *N* = 22), and large spheroids (mean 135.1 µm; range 99.7–158.7 µm; *N* = 27) analyzed for biomechanics in the extensional flow. **(e)** Minimum distance that the spheroid or cell center of mass approached the stagnation point *d_min_* normalized by the measured channel width *w*. This ratio reflected the degree to which the spheroid or cell felt the maximal hydrodynamic forces at stagnation point. Violin distribution plots: dot is the median, vertical line is the interquartile range (25th–75th percentile), horizontal line is the mean.

To compare the mechanics of the different sized spheroids and cells, device dimensions and infusion flow rates were design to achieve similar hydrodynamic extensional forces in the central cross-slot region. Devices were fabricated to have square cross-sections with a target channel-width-to-spheroid/cell-diameter ratio of three: *w/D*_0_ ∼ 3 (Supplementary Figure S2). Thus spheroids and cells experienced similar degrees of confinement and shear stresses in the straight channels [22] and similar wall effects from the channel corners in the extensional flow region. Cross-slot devices generate approximately planar extensional flow with a velocity field **v** = Ω(−*x***x̂** + *y***ŷ**) where Ω = −*∂u/∂x* = *∂v/∂y* is the extensional strain rate [23]. The characteristic extensional strain rate in the microfluidic devices was estimated as Ω ∼ Δ*u/*Δ*x* = (0 − *U*)*/*(*w/*2) = 2*U/w* where *U* is the average incoming fluid velocity in each straight inlet channel and *w* is the measured channel width. The flow rates for each experiment were selected to achieve approximately equal characteristic Ω values based on the target channel dimensions. However, small deviations in the fabricated cross-slot dimensions (channel width and depth) from the intended designs resulted in some variation in the realized extensional strain rates. The characteristic extensional strain rates for the single cell, small spheroid, and large spheroid experiments were Ω = 659, 779, and 896 s*^−^*^1^, respectively. The other parameter affecting the extensional hydrodynamic forces in the cross-slot is the buffer viscosity *µ*. The same buffer batch was used for all experimental groups to maintain consistent viscosity (zero-shear viscosity *µ* ∼ 90 mPa·s, Supplementary Figure S3). Thus the characteristic extensional forces *µ*Ω were 60, 71, and 82 Pa for the single cell, small spheroid, and large spheroid experiments, respectively. Although not identical, the device extensional forces of *µ*Ω = 71 ± 11 Pa were approximately matched across the three experimental groups with a reasonable variation of ±15%.

Spheroids and cells of different sizes showed similar speed trends and with some differences in deformation (**Figure 2A**, **Figure 2C**). Between groups, the speed values and dwell times varied due to the different flow rates and channel dimensions used for each group. To visually compare trends, time was normalized by the duration each cell/spheroid spent in the extensional flow (i.e., from entry to exit, corresponding to the region between the red and blue dashed lines). In the straight inlet channel, spheroids and cells approached the the extensional flow region at a constant inlet speed *V_in_* and constant initial shape, as seen in the flatter portions of the speed and deformation data before *t* = 0. The initial effective diameter *D*_0_ was estimated from the average equivalent diameter 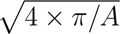 of the 2D projected areas *A* in the inlet channel (**Figure 2D**). The start of the extensional flow (vertical red dashed line in **Figure 2A**) was defined as the location in the microfluidic channel where the average cell or spheroid speed began to decrease from its constant *V_in_*. Our previous work showed that the start of extensional flow corresponded to where the velocity of incoming objects on the inlet channel centerline started decreasing and the end of extensional flow corresponded to the where velocity had recovered to a constant value in the outlet channel [23]. Under extensional fluid forces, the spheroids and cells stretched along the *y*-axis (outlet flow direction) and slowed down while approaching the flow stagnation point. Deformation was calculated from the 2D contour’s area moments of inertia along the *x* and *y* axes (more detail in Methods). The point of minimum velocity *V_min_* corresponded to the spheroid/cell’s minimum distance from the extensional flow stagnation point *d_min_*. The maximum deformation occurred near the end of the extensional flow region (blue dashed line, location where speed had risen back to a constant outlet speed) after these viscoelastic objects had sufficient time to deform under the hydrodynamic forces. Small and large spheroids displayed a larger change in inertia ratio compared to single cells (more below). All objects sped up to an exit speed in the straight outlet channel equivalent to the inlet speed (*V_out_ ≈ V_in_*).

The extensional hydrodynamic forces felt by spheroids and cells depended not only on the characteristic extensional strain rate of the unperturbed flow but also how closely it approached the stagnation point (*d_min_*). As shown from previous experimental [23] and numerical [24] analyses of the the unperturbed cross-slot fluid flow, within a distance about 20 - 33% of the channel width from the extensional flow stagnation point, the extensional strain rate was approximately uniform (Ω ∼ constant) and maximal. Further away, the extensional strain rate decreased quickly. Thus, *d_min_* normalized by the measured channel width *w* was used to assess how much cells and spheroids entered the constant extensional strain rate region (*d_min_/w <* 0.3) and served as a proxy to compare the hydrodynamic extensional forces felt by each experimental group. For all three groups, there was a range of *d_min_/w* values, and that range was roughly the same for the small and large spheroids (∼0.2 − 0.4) and overlapping although wider for the single cells (0.1 - 0.6) (**Figure 2E**). Taking into account the similar undisturbed fluid extensional forces (*µ*Ω = 71±11 Pa) and the similar variation in *d_min_/w*, the three experimental groups experienced similar extensional forces.

Though cells and spheroids felt similar extensional forces, it was more challenging for large and small spheroids to approach a cross-slot device’s central stagnation point compared to smaller single cells, even with upstream flow focusing to center the spheroids. The small single cells reached lower minimal speeds *V_min_* while the large spheroid had the largest *V_min_* values (**Figure 2C**, **Supplementary Figure S4**). Looking at all spheroids and single cells analyzed in the cross-slot, velocity data showed this trend of increasing *V_min_* with object size was true across experimental groups (**Supplementary Figures S4, S5a**). This result can be explained with irregular spheroid shapes, faster fluid velocities, and relative size of the cell or spheroid compared to the Ω ∼ constant region around the stagnation point. The irregular spheroid shapes that were not perfectly spherical meant that the fluid velocities and forces felt around the surface were not symmetric and a net force moved them towards one outlet. In contrast, single cell shapes were closer to idealized spherical and stretched ellipsoidal shapes with a smooth boundary. The higher flower rates and fluid velocities used for small and large spheroids more easily pulled the spheroids away from the stagnation point and towards the outlet compared to the smaller single cells. The finite sized objects in microfluidic extensional flow will have smaller *V_min_* values if the Ω ∼ constant region relatively large compared to the object because there is more time to dwell in the larger, low velocity region around the stagnation point [23]. Since the size of the Ω ∼ constant region scales with channel width [24, 23], the ratio of effective diameter to channel width *D*_0_*/w* can used to compare, in a relative sense, which group experienced the largest Ω ∼ constant region compared to its size. Here, the ratios of average diameter and channel width *D*_0_*/w* were 3.5 for single cells, 2.7 for small spheroids, and 2.0 for large spheroids. The target for all three groups was *D*_0_*/w* = 3, but device fabrication imperfections and larger than anticipated large spheroids led to differences. Thus, the larger *D*_0_*/w* value for single cells meant there was a relatively larger Ω ∼ constant region in which to dwell compared to the spheroids, leading to the increasing *V_min_* values.

### 2.2 Cell and Spheroid Deformation Observations

The spheroids showed a range of deformation behaviors in the extensional flow due to their irregular shapes and variation in cell-cell attachments. Four days of culturing resulted in spheroids without a necrotic core but also a range of shapes, including many with jagged edges. Some spheroids had slender protrusions that became highly stretched, bent, or rotated in ways that did not follow the bulk spheroid deformation (**Movie S4**). Rigid rotation in extensional flow was common due to the irregular spheroids having an effective long axis (e.g., of the best-fit ellipse to the 2D contour) that would be first oriented in the outflow flow direction before stretching (**Movie S5**). Due to the short times spheroids were in the cross-slot extensional flow region, misaligned spheroids would spend most to all of the extensional flow dwell time rigidly rotating with little to no stretching before leaving towards an outlet. Smaller spheroids frequently contained loosely attached cells and often lacked a strongly connected cellular core. As a result, many smaller spheroids displayed in- and -out-of-plane rotational motion in addition to deformation, while loose protrusions became highly stretched within the cross-slot region. In some cases, spheroids underwent substantial stretching and partial disintegration (**Movie S6**). These cases were typically associated with spheroids that were less spherical or contained loose cellular protrusions. Complete spheroid separation was rarely observed. More commonly, the loosely attached parts stretched significantly while remaining connected to the rest of the spheroid. In such cases, the spheroids generally did not recover their original shape in the outlet channel. Large spheroids sometimes recovered their original shape after exiting the extensional flow region, although recovery was often incomplete. In contrast, small spheroids exhibited greater variability in recovery behavior and frequently failed to recover their original shape, with some remaining substantially deformed in the outlet channel. We also observed occasional thin line-like structures extending from the spheroid, which were likely intracellular material released from a ruptured cell within the spheroid (**Movie S7**). These observations suggest that, in some cases, the internal stresses generated by collective spheroid deformation exceeded the mechanical tolerance of individual cells.

A subset of spheroids with nearly spherical morphologies that did not rotate were further analyzed, first from their 2D contour outline, similar to a common approach in single cell mechanical phenotyping [25, 26, 27, 28]. These quasi-spherical spheroids did not have loosely attached peripheral cells or prominent protrusions and also happened to be aligned with the outlet flow axis such that in extensional flow, these spheroids stretched without appreciable in-plane rotation. Spheroid deformation was quantified from the 2D contour with an area moments of inertia-based metric because the moments of inertia are known to be more robust to image noise and pixelation effects compared to aspect ratio, circularity and Taylor deformation parameter [29, 30] and reflect shape irregularities. The deformation was measured as the inertia ratio 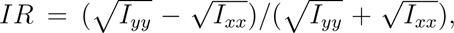 following previous analyses of droplet deformation in extensional flow [31, 32].

All near spherical, non-rotating spheroids and cells stretched due to the microfluidic extensional flow according to 2D contour deformation behavior, with cells stretching less than spheroids and small spheroids stretching more than large spheroids (**Supplementary Figure S4**, **Figure 2C**). The amount of stretching due to extensional flow was measured from the difference between the maximum and initial inertia ratios for each experimental group (**Figure S6a**). Single cells deformed less than spheroids under comparable extensional fluid forces. The change in inertia ratio is close to 0.1 in single cells and approximately 0.2 - 0.3 for both small and large spheroids, suggesting that single cells could be stiffer than the spheroids. This observation is consistent with findings of Panhwar et al. [18], who estimated that the Young’s moduli of spheroids (average diameter 134 µm, roughly the size of this study’s large spheroids) were approximately ten times smaller than that of single cells. Our results indicate that some smaller spheroids may be effectively softer than large spheroids. The change in inertia ratio between large and small spheroids was overlapping, but several small spheroids were highly stretched with stretching *>* 0.4 (**Supplementary Figure S4**). In many cases, the small spheroids unfolded into a nearly linear configuration with a large inertia ratio. These small spheroids exhibited relatively homogeneous deformation across their visible surface with many of the cells undergoing substantial stretching. In contrast, large spheroids maintained near spherical to ellipsoidal shapes with lower inertia ratios and consisted of a core region that had less visible deformation than the outermost cells. These observations were explored further with digital image correlation (upcoming Section 2.3).

When extensional flow ceased, spheroids traveling through the straight outlet channel often recovered towards their initial shape, though not consistently. Others retained a stretched in the outlet channel compared to their shape in the inlet channel, at least as measured from the 2D contour. Broadly speaking, small spheroids had greater variability in their recovery behaviors and tended to show less shape recovery compared to large spheroids, which partially recovered their initial quasi-spherical shape (**Supplementary Figure S4**). The variable recovery behaviors for small spheroids reflected their heterogeneous, irregular shapes. From observations, small spheroids had more possible shape conformations compared to quasi-spherical large spheroids because small spheroids consisted of only a few cells (typically 7–15 cells) that were tethered together at cell-cell junctions that made up a small fraction of individual cell surfaces. In contrast, the compact large spheroids had a sufficient number of cells such that some interior cells were completely surrounded by neighbors and those cells on the periphery were adhered to others through cell-cell junctions that covered a larger fraction of their surface area.

Overall, it was challenging to compare spheroid groups using deformations from 2D contours. The outline of the 2D projected spheroid shape does not capture the interior and surface stretching patterns of multicellular spheroids and out-of-plane stretching or rearrangements. The complicated and variable spheroid shapes along with a large portion of biomechanical information missing from 2D contour analysis meant that it was unsurprising that consistent deformation trends were not apparent using this approach. While 2D contour analysis has led to robust microfluidic mechanical phenotyping for single cells, the complex spheroids require a more detailed approach that considers deformations throughout more of the spheroid bulk material.

### 2.3 Digital Image Correlation Revealed Homogeneous Deformation in Small Spheroids and Core-Shell Heterogeneous Deformation in Large Spheroids

In microfluidic cross-slot stretching experiments, we observed that the outer layer of cells on large spheroids stretch significantly more than the central core, suggesting that the peripheral region may be mechanically softer than the inner core. To investigate this further, we applied digital image correlation (DIC) to stretched spheroids to measure the strain distribution. DIC is widely used in biomechanics to measure strain in tissues and bone, typically with speckle patterns applied to the sample [33, 34]. In spheroids under brightfield conditions, packed cells at slightly different *z*-levels generated sufficient pixel intensity texture for correlation analysis. DIC was applied to the subset of spheroids that did not rotate and had nearly spherical morphologies.

DIC analysis of spheroids deforming in extensional flow allowed for semi-quantitative comparison of small vs. large whole spheroid biomechanical behavior, though the experimental data was not ideal for high precision strain measurements. DIC is a 2D analysis technique for in-plane displacements of flat specimens, where here DIC was applied to 3D spheroids imaged from one overhead view. However, several features of our approach allowed for reasonable confidence in relative comparisons of DIC-measured strain (but not absolute values). Deformation by extensional flow limited out-of-plane spheroid strains that would reduce the accuracy of the 2D DIC analysis. Theoretically, an elastic sphere in planar extensional flow elongates into a prolate ellipsoid with the out-of-plane minor axis length remaining constant and unstretched [35]. The spheroids can be expected to deform similarly because they felt approximately planar extensional flow (the fluid velocities and extensional strain rates decreased closer to the top and bottom surface but still maintain the hyperbolic extensional flow streamlines). By limiting DIC analysis to only regions where spheroids felt extensional flow, the tumbling out-of-plane rotations that we observed in the straight inlet channel, particularly for small spheroids, was absent. Extensional flow, being an irrotational flow field, elongates and unfolds floppy objects, in contrast to shear flow that tumbles and balls up objects, as has been seen for single-stranded DNA [36, 37] and proteins [38]. Although there was volume averaging of DIC-measured strains due to the spheroid being partially transparent, our analysis comparing the core vs. periphery cells of its 2D projection was reasonable. For the partially-transparent spheroids viewed from below using an inverted microscope, images contained pixel intensity contributions from cells on the spheroid surface as well as interior cells that were above. So, DIC cross-correlation analysis would consist of a volume average (*z*-direction) of cells in a particular *xy*-plane neighborhood. However, given the quasi-spherical shape of the spheroids, cells on the periphery if the 2D projected contour did not have any cells above them, so the 2D DIC edge strains were dominated by only periphery shell cells. The 2D DIC interior strains would be dominated by the numerous 3D spheroid interior cells that were above. Finally, studies measuring intranuclear strains within living single cells [39] and spheroids [40] that were adhered to uniaxially stretching substrates demonstrate that DIC is possible in sub-millimeter samples.

DIC analysis was applied when spheroids were experiencing extensional flow, beginning when spheroids entered the extensional region in the inlet channel up to when extensional flow stopped in the outlet channel (**Figures 3A** and **4A**). The spheroid stretching movies were analyzed using the open-source 2D DIC software Ncorr [41] in MATLAB. Spheroids chosen for DIC analysis were selected based on spheroid morphologies that resulted in only elongational stretching and negligible rotation: minimal shape irregularities, no loose protrusions, and close to spherical shape. Over the time intervals in which spheroids experienced extensional flow, cumulative displacements were computed by adding together the displacements between consecutive frames while subtracting rigid motion, ensuring that only displacement due to deformation was retained. Green-Lagrangian strain components *E_xx_*, *E_xy_*, and *E_yy_* were estimated from the cumulative displacement using a local plane-fitting approach to compute displacement gradients. An equivalent 2D Von Mises strain *E_vm_* was calculated from these strain components. The DIC analysis approach was validated by digitally applying known deformations to spheroid images and comparing to NCorr DIC measurements. The validation tests for 10% strain, rigid body rotation and translation, and shear deformation showed that our DIC matched the digitally imposed deformations(Supporting Information Section 1.2 and Supplemental Figures S8–S11).

**Figure 3:**
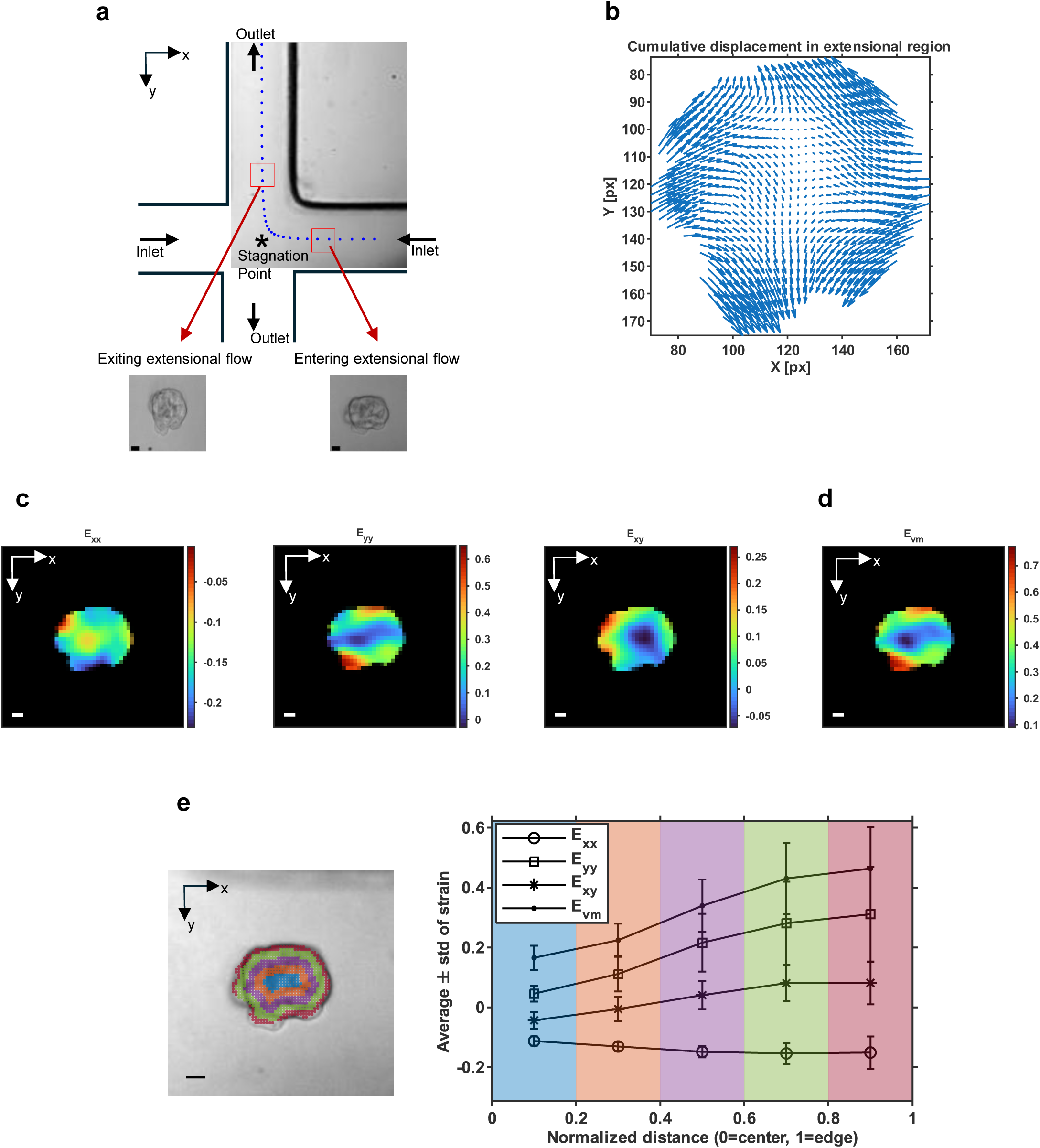
DIC spheroid strain analysis for a representative large spheroid (*D*_0_ = 99.7 µm). **(a)** DIC analysis was performed over the time interval in which the spheroid experienced extensional flow. **(b)** The cumulative displacement vectors. **(c)** Green-Lagrangian strain components *E_xx_* (inlet direction), *E_yy_* (outlet direction), and *E_xy_* (shear). **(d)** Equivalent von Mises strain (*E_vm_*). **(e)** Strain component mean ± standard deviation within spheroid radial layers. Scale bars 20 µm.

**Figure 4:**
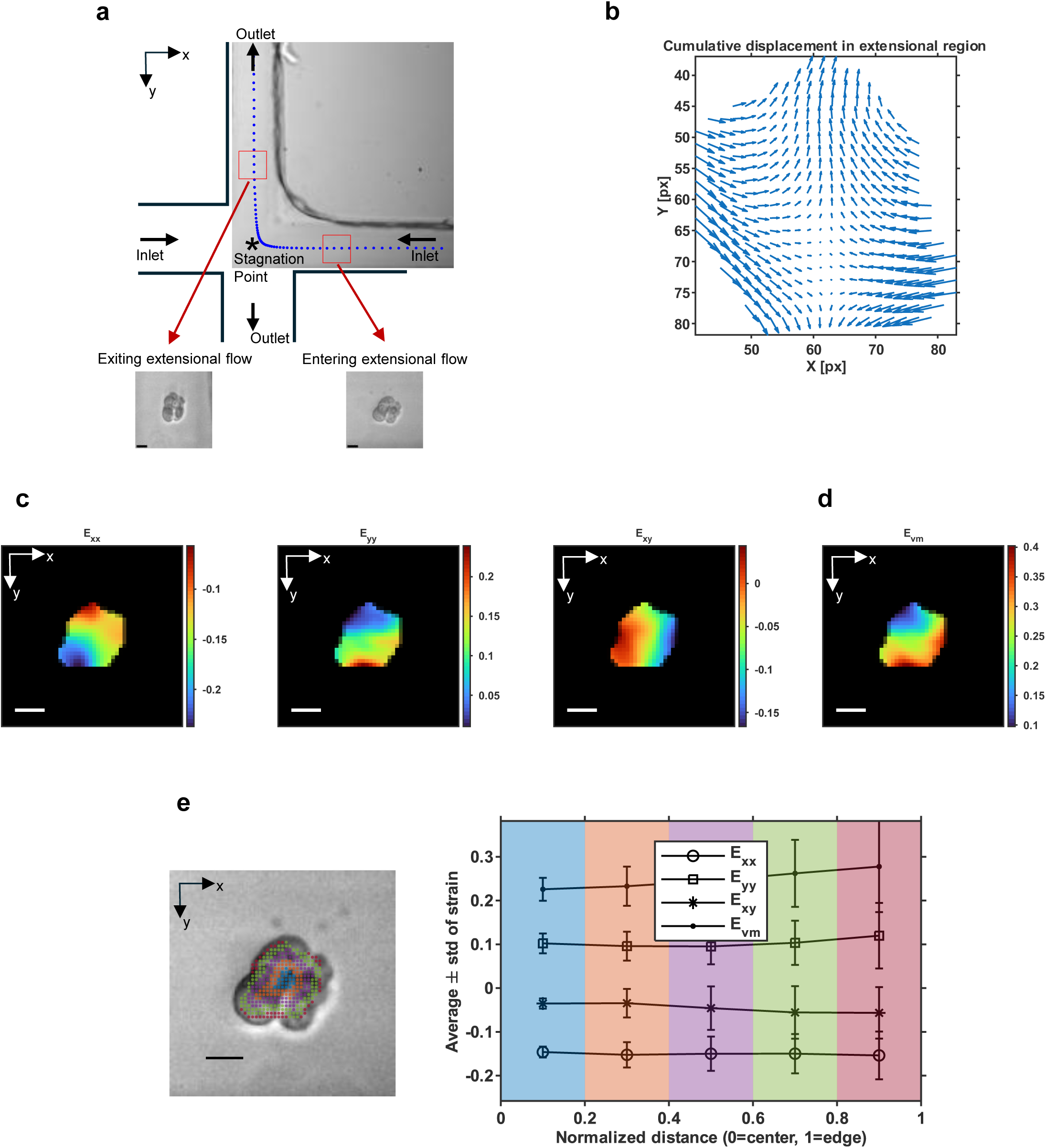
DIC spheroid strain analysis for a representative small spheroid (*D*_0_ = 47.5 µm). **(a)** DIC analysis was performed over the time interval in which the spheroid experienced extensional flow. **(b)** The cumulative displacement vectors. **(c)** Green-Lagrangian strain components *E_xx_* (inlet direction), *E_yy_* (outlet direction), and *E_xy_* (shear). **(d)** Equivalent von Mises strain (*E_vm_*). **(e)** Strain component mean ± standard deviation within spheroid radial layers. Scale bars 20 µm.

Overall, the DIC displacements and strains for large and small spheroids matched the theoretical exception that spheroids will deform into a prolate ellipsoids oriented with the outlet flow axis [35]. The DIC results for a representative large and small spheroid are shown in **Figure 3** and **Figure 4**. The cumulative displacement vectors showed that spheroids stretched along the outlet channel direction (*y*-axis) and contracted along the inlet channel direction (*x*-axis) (**Figures 3B** and **4B**). The spheroid elongation direction was also evident in the Green-Lagrangian strain components and equivalent von Mises strain *E_vm_*(**Figures 3C**, **3D**, **4C**, and **4D**). The *E_yy_* strain component was largest and dominated the equivalent von Mises strain. The *E_xx_* strains were negative due to compression in that direction, while the *E_xy_* shear strains were small throughout the spheroids.

To compare deformation trends throughout the representative large and small spheroids, Green-Lagrangian strains were compared across five concentric annular regions. For the large spheroid, the equivalent strain *E_vm_*and *E_yy_*both increased from the inner towards the outermost layer, suggesting that the outer layers stretched the most (**Figure 3E**). In addition, the positive *E_yy_* values meant that the large spheroid stretched along the *y*-axis. The shear strains *E_xy_* remained close to zero overall. *E_xx_* was approximately uniform and negative across the large spheroid, indicating compression along the *x*-axis. However, strain components and equivalent strain in the small spheroid were nearly uniform across all layers, despite locally elevated *E_vm_* at the outer edge (**Figure 4E**). This indicated that the small spheroid deformed relatively homogeneously, in contrast to the core-shell heterogeneity observed in the large spheroid. To further test our homogeneous small spheroid deformation conjecture, DIC analysis was performed on homogeneous and isotropic 0.5% w/w agarose microparticles stretching under extensional flow. Representative microparticle DIC results are shown in **Supplemental Figure S12**. The homogeneous agarose microparticle had uniform strain trends across the radial layers (**Supplemental Figure S12d**), closely matching the small spheroid strain trends and suggesting that small spheroids deform more homogeneously than large spheroids.

The contrast in inner and outer region deformation trends for small and large spheroids was consistent across many spheroids. Equivalent von Mises strain was compared between inner and outer regions for large spheroids (range *D*_0_ = 100–144 µm, *N* = 8), small spheroids (range *D*_0_ = 39–68 µm, *N* = 7) and agarose hydrogel microparticles, a homogeneous, isotropic control (range *D*_0_ = 31–50 µm, *N* = 8) (**Figure 5**). For this comparison, the inner region was defined as DIC subset centers with normalized radial distance of *r/D*_0_ < 0.25 from the spheroid center of mass, and an outer layer with *r/D*_0_ *>* 0.6 (**Figure 5A**). These thresholds were chosen to compare the obvious interior and peripheral spheroid regions while excluding the mixed intermediate region and also to ensure a sufficient number of DIC subset centers in each layer. The agarose microparticle comparison addressed whether outer region strains were elevated in all stretching objects regardless of composition, or whether the trend reflected genuine material inhomogeneity specific to large spheroids.

**Figure 5:**
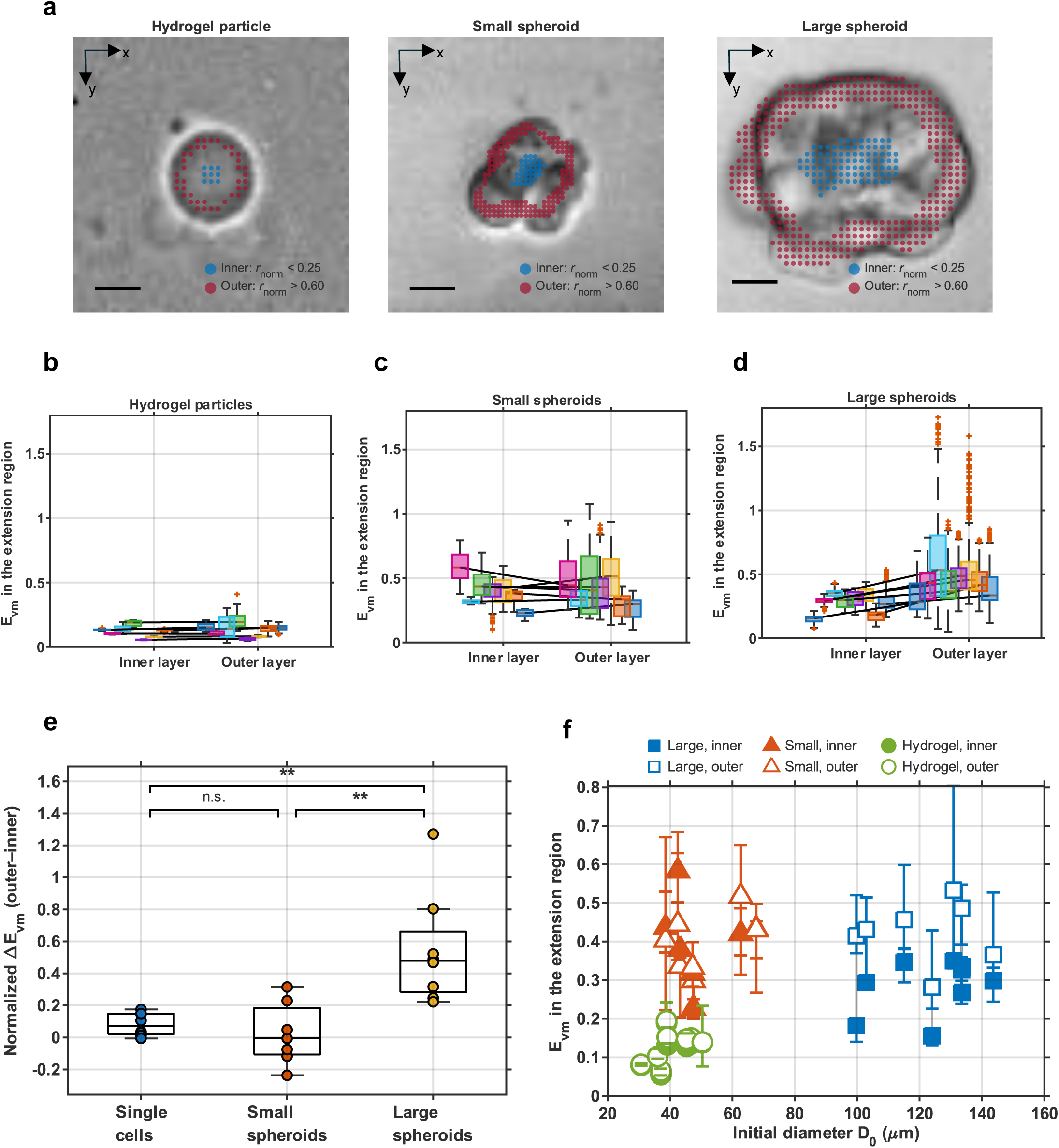
Comparison of DIC-measured inner and outer layer strains for large spheroids (mean *D*_0_ = 122.9 µm, range 100–144 µm, *N* = 8), small spheroids (mean *D*_0_ = 50.0 µm, range 39–68 µm, *N* = 7) and hydrogel microparticles (mean *D*_0_ = 40.6 µm, range 31–50 µm, *N* = 8). **(a)** Representative images of large and small spheroids and a hydrogel particle with the defined inner and outer layers. The inner layer corresponded to DIC subset centers with normalized distances from the particle center of *r/D*_0_ < 0.25 (blue dots), while the outer layer corresponded to *r/D*_0_ *>* 0.6 (red dots). Paired box plots of DIC equivalent von Mises strain distributions in the inner and outer layers for **(b)** agarose hydrogel microparticles, **(c)** small spheroids, and **(d)** large spheroids. Horizontal lines connect inner and outer box plots of the same color corresponding to a single spheroid. **(e)** Post hoc Dunn’s test pairwise comparisons of normalized von Mises strain median differences across the three experimental groups. Multiple comparisons were corrected using the Bonferroni method. Significant differences were observed between small and large spheroids (*p* = 0.003) and between large spheroids and hydrogel microparticles (*p* = 0.008), with no significant difference between small spheroids and microparticles. \*\**p <* 0.01 **(f)** Equivalent strain *E_vm_* versus initial particle diameter *D*_0_ for large spheroids (blue squares), small spheroids (orange triangles), and hydrogel particles (green circles) in the inner (filled) and outer open) layers. Markers show the median *E_vm_* within the layer and error bars span the interquartile range (25th–75th percentile).

Inner and outer layer von Mises strain distributions were visualized as box plots with horizontal lines connecting the median inner and outer strains for individual spheroids or microparticles. Across all groups, outer layer box plots generally exhibited larger spread, reflecting higher uncertainty in DIC measurements near particle boundary. The agarose microparticles showed tightly clustered, nearly identical inner and outer distributions, consistent with their homogeneous, isotropic composition; the inner-to-outer box plot connecting lines were correspondingly close to horizontal (**Figure 5B**). Like the microparticles, the small spheroids had similar inner and outer layer strain distributions, though with more variation due to the heterogeneous composition of biological cells compared to the simple agarose hydrogel material (**Figure 5C**). The contrast between inner and outer layer strains was the most pronounced for the large spheroids (**Figure 5D**). The median von Mises strain values increased from inner to outer regions for all large spheroids, and the outer layer had much higher strains and a wider distribution than the relatively tighter strain distributions in the inner layers.

The statistical significance of inner-to-outer layer differences across groups was tested on a normalized change in von Mises strain: Δ*E_vm_* = (med(*E_vm,out_*) − med(*E_vm,in_*))*/*med(*E_vm,in_*). A Kruskal Wallis test between the three experimental groups indicated that at least one group differed significantly in Δ*E_vm_* from the others (*p* = 0.001). Post-hoc Dunn’s tests to determine which specific groups differed showed significant differences between small and large spheroids (*p* = 0.003) and between agarose microparticles and large spheroids (*p* = 0.008), and no significant difference between small spheroids and hydrogel particles (**Figure 5E**). These results indicate that large spheroids deformed mechanically inho-mogeneously between their core and shell, while small spheroids deformed in a uniform manner similar to the isotropic hydrogel microparticles.

To examine whether this inner-outer divergence varies with particle size, we plotted the inner and outer layer median *E_vm_* strains and inter-quartile ranges against spheroid or microparticle initial diameter *D*_0_ (**Figure 5F**). This view complements the categorical comparison above by treating *D*_0_ as a continuous variable. Hydrogel microparticles showed closely overlapping inner and outer *E_vm_* strain distributions with negligible vertical separation between inner and outer med(*E_vm_*), regardless of particle size. Some small spheroids had larger med(*E_vm_*) in the inner region and others had a larger med(*E_vm_*) in the outer region. All small spheroids had substantial overlap in inner and outer strain distributions. All large spheroids had higher med(*E_vm_*) in the outer region compared to the inner region and much higher strains in the outer region.

Microfluidic extensional flow deformation of spheroids revealed interesting biomechanical differences between small spheroids (*D*_0_ ≲ 70 µm) and large spheroids (*D*_0_ ≳ 100 µm) with a cross-over point between ’large’ and ’small’ within the diameter range 70–100 µm, for which we have no spheroid biomechanical data in this work. Small spheroids tended to undergo significant stretching, with only partial and limited shape recovery after exiting the extensional flow region, consistent with a lack of mechanically dominant core (Figure 4). Large spheroids resisted deformation at their stiffer core while the outer shell deformed more strongly (Figure 3). Regional core-shell differences have also been reported in larger cancer spheroids (≥ 200 µm) under mechanical compression, where proliferation was preferentially inhibited in the interior [42]; although that study examined a biological rather than mechanical response, it supports the concept that distinct spheroid regions behave differently, consistent with the mechanical heterogeneity we observed in large spheroids.

Shape descriptors derived from the projected 2D contour, such as the inertia ratio based deformation used here, captured part of the overall spheroid deformation but could not resolve internal, spatially heterogeneous strain. In the end, the intra-spheroid DIC strain data provided the significant differences in small and large spheroids in this study. Due to the efficient deforming nature of extensional flow in stretching spheroids towards opposite outlets, the cross-slot extensional flow revealed heterogeneous strain within the large spheroids, distinguishing deformation of the core from that of the outer shell. For small spheroids with cell-cell connections covering less of the cells’ surfaces, the extensional flow unfurled the folded cell clusters to expose the stronger cell-cell junctions. In contrast, spheroids in the shear-dominated parabolic flow of a straight microfluidic channel will deform into a relatively compact bullet-like shape [18], making it potentially more challenging to identify spheroid mechanical and structural heterogeneities.

### 2.4 Cell Circularity and Nuclear Sphericity Decreased with Increasing Spheroid Size

To begin to explain the differences in large and small spheroid deformation under extensional flow, we investigated cell morphology within the spheroids. From visual observations, most large spheroids exhibited a distinct core-shell structure, with a dense central core of packed and interconnected cells surrounded by a shell of more loosely connected peripheral cells. Smaller spheroids typically lacked this clearly defined core. In previous studies of organoids embedded in an extra cellular matrix, interior core cells had smaller nuclear volumes than peripheral cells, an effect attributed to compression-driven fluid movement through cell-cell junctions; the resulting swollen peripheral cells were softer and more metastatic [21, 43]. Furthermore, changes in cell volume are known to alter cellular stiffness, with cells becoming stiffer as they shrink and softer as they swell [44, 45, 46]. Volume changes also modify cytoskeletal and focal adhesion formation and the density of intracellular proteins [47], all of which can affect individual cell mechanics and bulk spheroid deformation.

To determine whether large PANC-1 spheroids showed a similar core-shell difference in cell/nuclear volume that could related to their mechanical heterogeneity, live spheroids were stained with plasma membrane, F-actin, and nuclei dyes and imaged with confocal microscopy (**Figure 6A** and **6B**). To mimic spheroid shape in suspension, short incubation times were used to minimize spheroid adhesion to and spreading over the glass substrate. In general, cells within small spheroids maintained a round shape and appeared uncompressed. In contrast, cells within large spheroids were more densely packed; cells in the inner region were smaller and outer cells were often larger.

**Figure 6:**
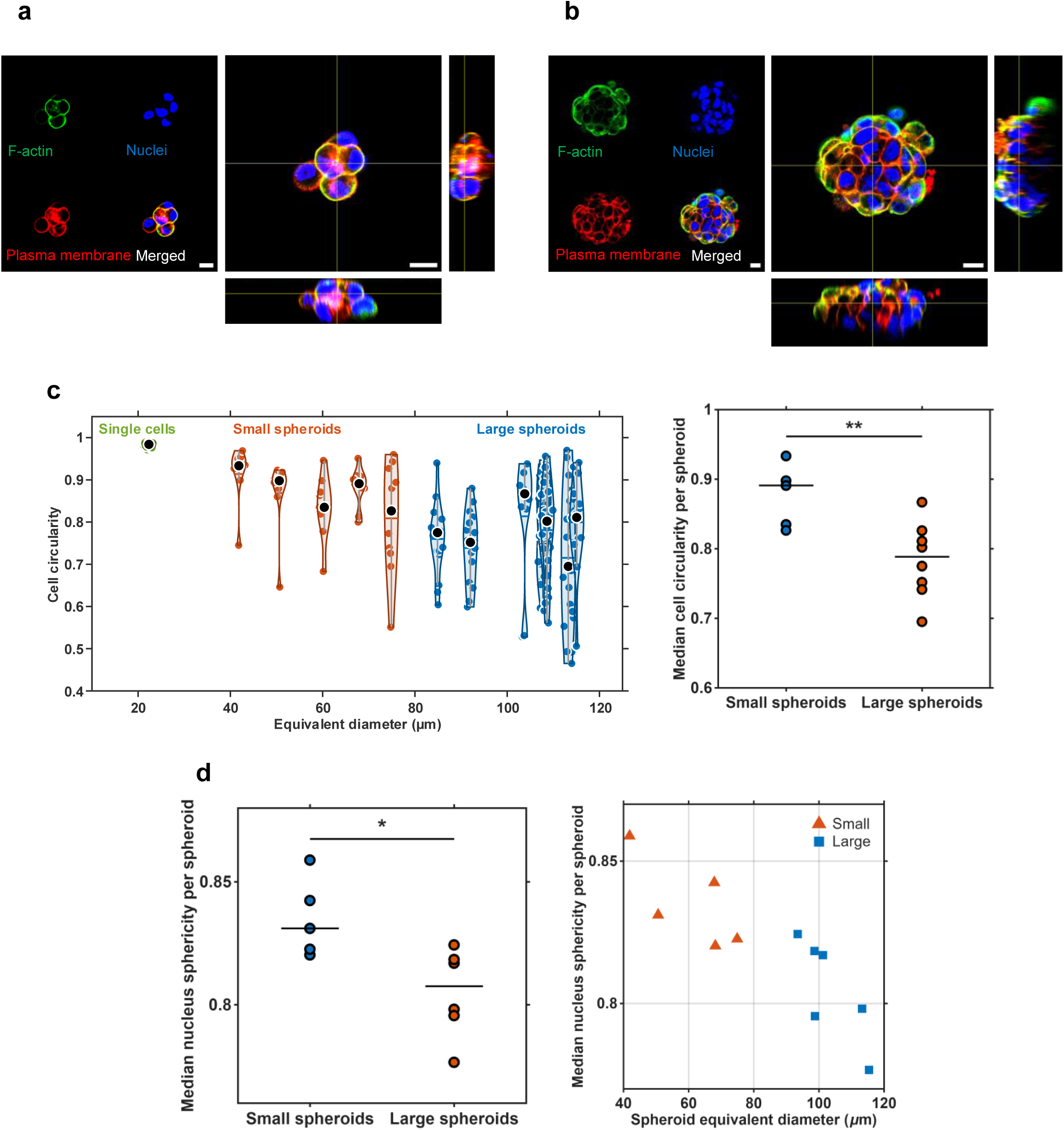
Morphology analysis of individual cells within small and large spheroids. Representative confocal midheight and orthogonal view images of **(a)** a small spheroid (*D*_0_ = 50.6 µm), and **(b)** a large spheroid (*D*_0_ = 113 µm) with live-cell staining. Scale bars are 20 µm. **(c)** 2D cell circularity analysis from the stained plasma membrane for cells at the spheroid midheight. Left: Cell circularity distributions within spheroids compared to single cells. Blue shaded region: suspended single cells (*N* = 19) under brightfield imaging conditions. Unshaded region: circularity of individual cells within small spheroids (mean *D*_0_ = 159.1 µm, range 41.8–74.9 µm, *N* = 5) and large spheroids (mean *D*_0_ = 104.2 µm, range 84.9–115.1 µm, *N* = 8). The red dashed line separates small and large spheroids defined in this section. Violin plots: Black dot: median, horizontal line: mean. Right: Median differences of 2D cell circularity between small and large spheroids were assessed using a Mann-Whitney U (rank-sum) test (*p* = 0.006). **(d)** Median 3D nuclear sphericity for cells throughout the spheroid volume. Left: Comparison of median nucleus sphericity between small (mean *D*_0_ = 60.7 µm, range 41.8–74.9 µm, *N* = 5) and large spheroids (mean *D*_0_ = 103.5 µm, range 93.5–115.4 µm, *N* = 6). Differences were assessed using a Mann-Whitney U (rank-sum) test (*p* = 0.017). Right: Median nuclear sphericity decreased with spheroid size as cells became more packed in the inner core region. The red dashed line separates small and large spheroids. \**p <* 0.05 \*\**p <* 0.01

Cell shape and packing was first assessed from the 2D circularity of cells near the mid-height equatorial plane. For this analysis, cell boundaries were determined from plasma membrane images. When comparing the distributions of cell circularity, cells within all spheroids were substantially less circular than suspended single cells (**Figure 6C**). Cell circularity distributions showed greater variability and lower medians in large spheroids (range *D*_0_ = 85–115 µm, *N* = 8) compared to small spheroids (range *D*_0_ = 42–75 µm, *N* = 5), indicating an increased proportion of compressed, less circular cells with increasing spheroid size (Mass-Whitney U test, *p* = 0.006). A similar comparison of median cell area per spheroid for cells near the spheroid mid-height showed decreased median cell area for large spheroids compared to small, but not at a statistically significant level (*p* = 0.093, Mann-Whitney U test, **Supplemental Figure S13a**).

Three-dimensional cell size and shape was characterized using nuclear shape as proxy [43, 21]. Median nuclear sphericity, where 1 indicates a perfect sphere, was significantly lower in large spheroids (range *D*_0_ = 95–115 µm, *N* = 6) than small spheroids (range *D*_0_ = 42–75 µm, *N* = 5) according to the Mann-Whitney U test, (*p* = 0.017) and decreased with increasing spheroids diameter (**Figure 6D**). Expecting that packed interior cells would have smaller volumes than those in outer layers, we compared nuclear volume between the core and shell regions within small and large spheroids. However, neither spheroid size group showed a significant difference between core and shell median nuclear volume, with *p*-values of 0.31 for large spheroids and 0.44 for small spheroids using a Wilcoxon signed-rank test (**Supplemental Figure S13b**).

Taking together the 2D cell circularity and 3D nuclear sphericity results, there is some evidence that large spheroids (*D*_0_ ≳ 95 µm) possessed a packed core of cells with looser packing of the peripheral cells while small spheroids (*D*_0_ ≲ 75 µm) consisted of closer-to-spherical cells adhering together. However, the contrasts between the core and shell regions for our spheroids in a quasi-suspended state were not as pronounced as for embedded spheroids. Long et al. [43] observed pronounced differences in nuclear volumes between core and periphery of ECM-embedded organoids. Upon release from the ECM and relaxation in a stress-free environment, core nuclear volumes increased while peripheral nuclear volumes decreased slightly, reducing the disparity between regions. Consistent with this, our spheroids, imaged live and unembedded, and so never subject to ECM confinement, showed no significant difference in nuclear volume between core and shell. Our nuclear volume measurements were also consistent in magnitude with PANC-1 nuclear areas reported by Nguyen et al. [48] (converted to an equivalent spherical volume of 1196–3642 µm^3^) despite our measurements being taken from cells within intact spheroids rather than dissociated single cells.

Changes in cell volume resulting from mechanical or environmental cues can directly influence biochemical pathways and, consequently, cell fate. A reduction in cell volume increases macromolecular crowding by confining cellular components in a smaller space, which can promote biomolecular phase separation and modulate Wnt/*β*-catenin signaling; depending on tissue context, this can either enhance or suppress proliferation [49, 50]. In addition, changes in cell volume more broadly increase cell-to-cell variability in biochemical processes even within an initially homogeneous population [51]. Although the differences in 2D cell area and 3D nuclear volume we observed across spheroid sizes and within intra-spheroid layers did not reach statistical significance, cells in larger spheroids trended toward smaller and more variable areas and became significantly less circular and more heterogeneous in shape, suggesting that spheroid growth may introduce cell-to-cell variability that could, in turn, diversify biochemical states within the population.

### 2.5 Vimentin Showed Shell-Enriched Expression in Large Spheroids and Homogeneous Expression in Small Spheroids

Given the core-shell mechanical and cell shape heterogeneity in large spheroids, we hypothesized that this pattern may partly reflect differential protein expression between the core and shell. Spheroids of various sizes were fixed and immunostained for vimentin, E-cadherin, and (in additional samples) F-actin, with nuclei labeled by DAPI. Vimentin is a cytoskeletal intermediate filament protein that maintains cell structure and stiffness, protects the nucleus under compression, and contributes to cell migration and invasion [52, 53, 54, 55, 56]. E-cadherin, an adherens-junctions protein linked to the actin cytoskeleton, maintains cell-cell adhesion and responds to tensile mechanical cues [57, 58, 59, 60]. High vimentin combined with low E-cadherin expression is a canonical marker of epithelial-mesenchymal transition (EMT) and increased metastatic potential.

Expression of vimentin was relatively uniform throughout small spheroids while large spheroids primarily expressed vimentin in the outer cell layer. In small spheroids (range *D*_0_ = 61–72 µm, *N* = 4), vimentin was uniformly expressed around nuclei throughout the spheroid (**Figure 7A**). In large spheroids (range *D*_0_ = 76–115 µm, *N* = 7), vimentin was absent around core nuclei while the outer shell retained substantial vimentin expression (**Figure 7B**). For all spheroids, there were less apparent trends in E-cadherin, F-actin, and nuclear stain levels throughout the spheroids. visually looked pretty UNIFORM

**Figure 7:**
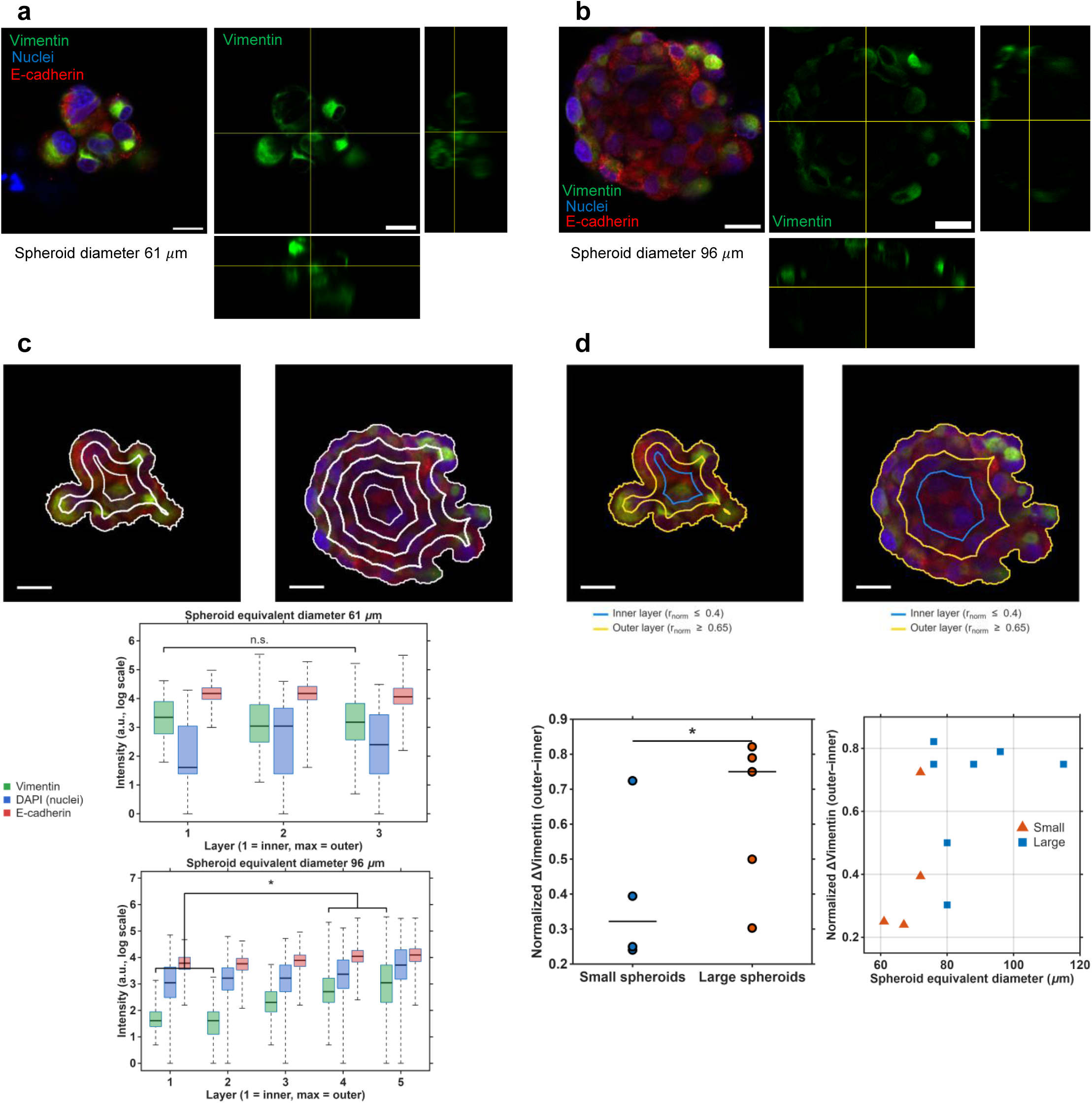
Vimentin, E-cadherin, and nuclei distributions within small and large spheroids from 3D confocal immunofluorescence imaging. All scale bars are 20 µm. Spheroid mid-depth and orthogonal views (vimentin only) of **(a)** a representative small spheroid (*D*_0_ = 61 µm) and **(b)** a representative large spheroid (*D*_0_ = 96 µm). **(c)** Analysis of the distribution of vimentin and E-cadherin and nuclei (DAPI) intensities in 8-bit images within concentric layers of representative small (left) and large (right) spheroids. The vertical axis is log(intensity + 1). The 10-µm-wide layers (white outlines) were roughly the thickness of a layer of cell. The difference in vimentin intensity between two innermost and two outermost layers was significant in large spheroids using the Wilcoxon signed-rank test (*p* = 0.016). In contrast, small spheroids did not show a significant difference in vimentin intensity between the innermost and outermost layers (*p* = 0.125). The same test showed no significant difference in E-cadherin intensity in either small (*p* = 0.25) or large (*p* = 0.078) spheroids, or in DAPI intensity in small spheroids (*p* = 0.75). DAPI intensity in large spheroids differed significantly according to the *p* ¡ 0.05 threshold, though marginally (*p* = 0.047).**(d)** Statistical comparisons between the inner (red shaded region) and outer (blue shaded region) regions overlaid for small and large spheroids. The inner layer was defined as normalized radial distance *r/D*_0_ < 0.4 from spheroid center, and the outer layer was normalized radial distance *r/D*_0_ *>* 0.65. Mann-Whitney U test between small spheroids (mean *D*_0_ = 67.9 µm, range 60.8–72.2 µm, *N* = 4) and large spheroids (mean *D*_0_ = 87.3 µm, range 75.8–115 µm, *N* = 7) showed a significant difference in the normalized vimentin difference between the inner and outer layers (*p* = 0.036). The normalized vimentin difference was defined as the difference between inner and outer layer vimentin median intensities normalized by the outer layer vimentin median intensity.

The spatial variation in protein expression levels within spheroids was further assessed by comparing data from concentric layers (**Figure 7C**, **Supplemental Figure S14a**). For small spheroids, image intensity levels for DAPI (nuclei), vimentin, and E-cadherin were approximately uniform across layers, and a Wilcoxon signed-rank test comparing the inner and outer layer vimentin intensity was not significant (*N* = 4, *p* = 0.125). For large spheroids, nuclei and E-cadherin were approximately uniform across layers, but vimentin was substantially lower in the inner two layers and fully expressed in the outer two layers, a significant inner-outer difference (*N* = 6, *p* = 0.016). Given that the fluorescent signals from the DAPI nuclei stains were consistent throughout the large spheroid volumes, the low vimentin fluorescent single levels in the interior of large spheroids was likely due to a lack of vimentin in the interior as opposed to signal attenuation due to tissue absorption.

To quantify the vimentin expression patterns further, we defined a normalized vimentin difference between in the inner (*r/D*_0_ < 0.4) and outer (*r/D*_0_ *>* 0.65) spheroid regions: ΔVim = (med(Vim*_out_*)–med(Vim*_in_*))/med(Vim*_out_*). This normalized inner-to-outer vimentin difference was significantly higher in large spheroids (range *D*_0_ = 76–115 µm, *N* = 7) than in small spheroids (range *D*_0_ = 61–72 µm, *N* = 4) according to the Mann-Whitney U test (*p* = 0.036, **Figure 7D**). The same comparison for DAPI and E-cadherin showed no significant differences (*p* = 0.412 and 0.073, respectively). This shell-enriched vimentin pattern in large spheroids is consistent with Cramer et al. [61], who reported vimentin expression restricted to the outer invading cells of ECM-embedded PANC-1 spheroids.

The uniform E-cadherin and F-actin levels were reasonable for the PANC1 cell line used in this study. Here, E-cadherin expression did not differ substantially among spheroid sizes and remained relatively uniform throughout each spheroid regardless of vimentin level, rather than following the general inverse vimentin–E-cadherin EMT relationship found in many cell lines. PANC-1 is known as a quasi-mesenchymal line with low E-cadherin and high vimentin mRNA expression [62, 63, 64]. Our results reproduced this stable low baseline of E-cadherin expression in PANC-1 spheroids. F-actin expression remained consistent across spheroid sizes (**Supplemental Figure S14b**). F-actin and vimentin form an integrated cortical network rather than acting independently [65]. For this study, the constant F-actin and variable vimentin distributions for large spheroids indicated that, if cell cytoskeletal mechanics differences impacted spheroid bulk deformation, then vimentin was likely a larger contributor to the observed core-shell mechanical behavior than F-actin.

### 2.6 Discussion

Using the cross-slot microfluidic technique, we showed that large spheroids (average *D*_0_ *>* 90 µm) exhibit core-shell heterogeneous mechanical behavior, in contrast to small spheroids (average *D*_0 ≈_ 50 µm), which deform homogeneously. We linked this heterogeneity in large spheroids to vimentin expression in the shell, and the sphericity of cells within the spheroids. Though this is a preliminary study with relatively small numbers of spheroids in the analyses, the results proposed a cross-over region that could be studied more in future work. The presence of mixed ΔVim results in intermediate sizes, compared to a monotonic trend, along with statistically significant small vs. large spheroid comparisons across all three data sets suggested a transition region of spheroid size of 75 < *D*_0_ < 90 µm that lies between the small and large spheroid groups with different characteristics for biomechanics, cell morphology, and vimentin cytoskeletal protein expression.

A recent study demonstrated that multicellular spheroids can transition between distinct mechanical states through changes in cellular organization and biochemical regulation. Marchesi et al. [66] showed that molecular regulators promoted tissue fluidization and collective spreading by altering cell-cell interactions, cellular rearrangements, and effective tissue viscosity. Although the present study examines passive deformation under externally applied extensional flow rather than active spreading, our findings similarly emphasize the importance of tissue-scale organization in governing spheroid mechanics. Together, these studies support the view that the mechanical behavior of multicellular tumor spheroids arises from a complex interplay between cellular composition, structural organization, and biochemical regulation. Understanding these relationships will be important for the development of physiologically relevant spheroid models and for elucidating the role of mechanics in tumor progression.

Two of the key parameters known to influence spheroid stiffness are the stiffness of individual cells composing the aggregate and the strength of the cell-cell interactions between them. Computational modeling of micropipette aspiration has shown that cell-cell adhesion, rather than single-cell stiffness, dominates a spheroid’s bulk elastic modulus, although both factors influence the timescale of its viscoelastic response [67]. Based on our observations from deformation experiments with both single cells and spheroids, we hypothesize that single cells change their mechanical properties once they become part of an aggregate. These changes may depend on the number of cell-cell junctions (size of the spheroid), as well as the cell’s location within the spheroid. Single cells in suspension appear spherical, whereas once incorporated into an aggregate they adopt a generally more deformed, non-spherical shape. Vimentin is known to be expressed in suspended single cells [68], but in large spheroids, cells in the core and shell expressed different vimentin intensities. A similar stiffer core and softer periphery organization has been predicted by computational models of 3D tumor growth [69]. These observations raise further questions about how cell stiffness is modulated by cell-cell contact. For example, does the softness of outer-layer cells in larger spheroids correspond to an increased propensity for metastasis, given that reduced cell stiffness has been linked to greater metastatic and invasive potential in other cancer types [70, 71]? As a next step, it would be informative to treat spheroids with drugs that target cell-cell junctions to determine how disrupting these junctions alter overall spheroid stiffness. Combining our extensional flow approach with a complementary technique capable of resolving the stiffness of individual cells within an aggregate would also be a valuable direction for future work.

It is important to note that the core-shell mechanical behavior reported here was observed in PANC-1 spheroids, which originate from a quasi-mesenchymal pancreatic ductal adenocarcinoma cell line[72]. PANC-1 spheroids typically exhibit irregular morphology, loose cellular organization, high vimentin expression, and low E-cadherin expression, resulting in less compact aggregates than those formed by more epithelial pancreatic cancer cell lines [62, 63, 64]. Consequently, the mechanical responses observed in this study may differ from those of spheroids formed from other PDAC cell lines that retain more epithelial traits. PDAC cell lines with a more epithelial phenotype. Future studies should investigate whether similar mechanical heterogeneity is present in other pancreatic cancer cell lines, including the more chemoresistant PANC1-OR subline [73], as well as in other mesenchymal and quasi-mesenchymal cancer cell lines.

## 3 Conclusion

We demonstrated that PANC-1 spheroid size strongly influenced mechanical behavior during stretching in microfluidic extensional flow. Small spheroids (mostly *D*_0_ = 40–75 µm) deformed relatively homogeneously and uniformly, with no evidence of a distinct bulk core, allowing substantial deformation under extensional flow. In contrast, larger spheroids (mostly *D*_0_ = 90–145 µm) exhibited a heterogeneous core-shell mechanical response characterized by a stiffer core and a softer, more deformable shell. Analysis of cellular morphology, together with vimentin expression, further revealed regional core-shell differences within large spheroids that were not observed in smaller spheroids. There was a transition size region between large and small spheroids that was roughly *D*_0_ = 75–90 µm. Where our study had data in this intermediate spheroid size region, vimentin protein expression trends were mixed while there was a more consistent small spheroids and large spheroid features and trends on either side. Additional experiments in this cross-over spheroid size region could further clarify the boundaries and patterns in each region.

Our findings highlight the importance of spheroid size in determining mechanical behavior. As spheroids mature, changes in cell number, cell-cell interactions, and cellular packing collectively alter their mechanical properties. Because mechanical properties are closely linked to underlying biochemical and structural characteristics, our results suggest that spheroids of different sizes may also possess distinct biological phenotypes. This consideration is particularly important for tissue engineering applications and for the use of spheroids as *in vitro* cancer models. Understanding the relationship between mechanical behavior and biological properties in cancer is essential for developing more effective diagnostic and therapeutic strategies [4, 5]. In this study, we examined a limited set of biological markers to investigate possible molecular mechanisms that explained the large spheroid mechanical heterogeneity. Additional immunofluorescence studies could provide further insights into the observed mechanical heterogeneity. In addition to vimentin and E-cadherin, proteins involved in adherens and tight junctions, such as *β*-catenin and ZO-1, may help elucidate the cell-cell interaction mechanisms governing tissue integrity and mechanical behavior [58].

Microfluidic extensional flow complements the suite of existing mechanical testing approaches for spheroids, organoids and microscale tissue samples. Extensional flow has the advantage of opening up compact, globular objects into a more linear shape. This makes weak internal connections readily apparent and intra-spheroid deformation heterogeneity more visible compared to shear flow. The advantages of extensional flow for applications in tissue engineering could draw inspiration from single-molecule DNA manipulation [37, 74, 75, 76] and protein bioprocessing applications [38, 77] in which the uniform stretching has facilitated efficient analysis of internal sequence and structural architecture and controlled protein and/or DNA interactions and manipulations.

## 4 Materials and Methods

### Cell Culture

PANC-1 cells (ATCC) were cultured in 75 cm^2^ T-flasks (BioLite^TM^, Thermo Fisher) at 37 °C in a humidified incubator with 5% CO_2_ and 95% air. Cells were maintained in high glucose Dul-becco’s Modified Eagle Medium (DMEM; HyClone) supplemented with 10% fetal bovine serum (Gibco), 100 µg/mL Penicillin-Streptomycin (HyClone), and 0.5 µg/mL Amphotericin B (Corning). PANC-1 cells were passaged upon reaching 60–80% confluency, approximately once a week.

### Spheroid Preparation

PANC-1 spheroids were formed as non-adherent spheroids on an agarose menis-cus in media supplemented with dilute Matrigel, following the method previously reported by Cramer et al. [61]. To produce the large number of spheroids required for microfluidic flow experiments, we used a high-throughput adaptation of this method based on 256-well 3D Petri Dish micro-molds cast in agarose (Microtissues, Inc.). For sufficient numbers of spheroids for microfluidic experiments, twelve non-adherent 1.5–2 % w/w agarose Petri Dishes were prepared, placed in a 12-well plate, and equilibrated with media twice for 15 minutes each in the incubator before seeding single cell suspensions. Different spheroid sizes were obtained by adjusting the cell seeding density while keeping the 4-day growth period constant: ∼2 cells per micro-well to target 50 µm diameter spheroids and ∼15 cells per micro-well for 100 µm diameter spheroids. 120 µL of cell suspension was added to each agarose Petri Dish, followed by 10 minutes of cell settling time and the addition of 1.5 mL of media. Twenty four hours after cell seeding, 1.5 mL of 1 % v/v Matrigel (Corning, Cat. No. CB-40230C) in DMEM was added to each Petri Dish. These added extracellular matrix components promoted more spherical shapes and improved spheroid toughness for transferring steps. Spheroids were cultured for 4 days to allow for a cohesive spherical cell cluster structure while limiting the likelihood of a sizable necrotic core (Figure 1B). Spheroids were harvested by inverting the agarose Petri Dishes into 1 mL of warm media and gently agitating. Collected spheroids were gently centrifuged to obtain a spheroid pellet for subsequent use in microfluidic and imaging experiments.

### Spheroid Size and Viability Measurements

Spheroids were imaged in the 256-well micro-molds to assess spheroid growth and necrotic core development over two weeks. For this experiment, the agarose Petri Dishes were placed in 6-well glass-bottom plates (MatTek), which allowed the spheroids to be imaged in situ within the agarose Petri Dishes. The agarose Petri Dishes fit directly over the coverslip inset area of the MatTek plates. At each imaging time point (days 1–15, excluding days 6, 9, and 11), an average of 15 individual spheroids were analyzed for diameter and 15 spheroids were measured for viability. Equivalent spheroid diameters were determined from binarized dark-field images acquired using a 5× objective. Spheroids were segmented from the images, and their projected areas were used to calculate equivalent diameters assuming a circular shape. Spheroids were stained with Calcein AM (Corning Life Sciences) and ethidium bromide to identify live and dead cells, respectively. Spheroids were imaged using a 5× objective with a Zeiss AxioCam HRm camera mounted on a Zeiss AxioObserver.Z1 microscope. To quantify the necrotic area, raw .czi images corresponding to the dead-cell indicator (red fluorescence) were binarized in MATLAB, and the dead-cell area was quantified using the `regionprops` function, following previously described methods [78]. The spheroid size and viability measurements were repeated in four independent experiments (biological replicates).

### Microfluidic Device

All microfluidic devices used in the three experimental groups (single cell, small and large spheroids) were cross-slot devices (Figure S2). Each device consisted of two straight perpendicular channels intersecting in a cross to produce an extensional flow flow with zero velocity at the central stagnation point. To create similar hydrodynamic force conditions for the different sized spheroids and single cells, devices were fabricated to have square cross-sections with a target ratio of channel width to spheroid diameter equal to three, as the distance of microparticles from channel walls along with their speed influences the viscous shear and extensional forces applied. Additionally, the straight channel length between the flow focusing region and the cross-slot in all devices was set to 2 mm to ensure that the flow reached a steady state before entering the cross-slot. The microfluidic devices were fabricated using standard soft lithography techniques. Device masters were made by laminating layers of dry-film photoresist (Riston GoldMaster 50 µm-thick GM120 and 100 µm-thick GM140, DuPoint) onto stainless steel wafers (Stainless Steel Supply). The channel designs were transferred onto the wafers with a Karl Suss MJB3 contact mask aligner, followed by development in 10% potassium carbonate solution. Poly-dimethylsiloxane (PDMS) was cast onto the masters and cured at 60 °C overnight. Final PDMS devices were bonded to glass slides using a plasma cleaner (Herrick). The fabricated device channel dimensions differed slightly from the targeted dimensions. For the ∼ 50 µm spheroids, the intended 150 µm square cross-section channel was fabricated with a width of 150 µm and depth of 137 µm. For the ∼ 100 µm spheroids, the intended 300 µm square channel was fabricated with a width of 264 µm and depth of 268 µm. For ∼ 20 µm single cells, the desired 60 µm channel was fabricated with a width of 70 µm and depth of 55 µm.

### Aqueous Methyl Cellulose in PBS Solution

A 0.8 % w/v methyl cellulose solution in DPBS (1×) containing calcium (Gibco, Thermo Fisher Scientific) was prepared by dissolving methyl cellulose (Thermo Scientific Chemicals, CAS 9004-67-5, molecular weight 454.513 g/mol) in DPBS. Calcium was necessary to maintain the integrity of calcium-dependent adherens junctions between cells. The solution was vacuum filtered through a 2.5 µm paper filter and stored at 4 *^◦^*C. Viscosity was measured using a TA Instruments Discovery Hybrid series rheometer with a 40 mm diameter parallel plate geometry. This highly viscous buffer solution (viscosity ∼ 90 mPa·s) meant many spheroids were observed near the device channel mid-height and did not settle due to gravity in the syringe or in the device. More information about viscosity measurement is discussed in the Supplementary information.

### Spheroid and Single Cell Microfluidic Extensional Flow Experiments

Prior to microfluidic experiments, the spheroid pellet obtained after harvesting was gently resuspended in 8 mL (large spheroids) or 6 mL (small spheroids) of aqueous methyl cellulose and loaded into a syringe. Single cells were suspended in aqueous methyl cellulose at a concentration of 200,000 cells/mL. All experiments were performed under laminar flow conditions (Re < 1). For each experiment, the suspended sample and blank aqueous methyl cellulose were infused into the device with syringe pumps (Pump33 DDS and Pump11 Elite, Harvard Apparatus). The sheath-to-sample flow rate ratio was set to one to confine most spheroids and cells near center in the channel width direction. The total flow rate (sheath and sample) entering each side of the cross-slot region was 324 µL/hr for single cells, 4.3 mL/hr for small spheroids, and 30 mL/hr for large spheroids. Spheroids and single cells were observed with brightfield microscopy (Zeiss Axiovert 200M inverted microscope, 10× objective, 0.9983 pixels/ µm). Movies were acquired with a high-speed camera (JetCam 19M with Komodo Frame Grabber, Kara Instruments) and StreamPix 8 software (Norpix Inc.) with frames per second and exposure times of 2200 fps and 2.126 µs for single cells, 3000 fps and 1.6 µs for small spheroids, and 3200 fps and 1.885 µs for large spheroids. For efficient movie capture of single spheroids and cells stretching in extensional flow, the StreamPix Motion Detection module with Pre/Post triggering. The camera region of interest (ROI) included part of the sample inlet channel, the cross-slot region, and a portion of one outlet channel, with a bias towards the outlet channel to observe spheroid recovery after leaving the extensional flow region. Movies were recorded at least 1 minute after flow was initiated at a specific rate to ensure steady-state conditions. The microscope stage *z*-height was chosen so that the interior of cells and spheroids appeared slightly darker with a bright halo. This imaging approach facilitated subsequent analysis for tracking and contour detection, as suggested by Herbig et al [30].

### Spheroid Deformation Analysis from 2D Contours

Streampix sequence files were directly imported to MATLAB by using the ReadJpegSEQ [79]. Approximate spheroid/cell shapes and locations in each movie frame were identified from pre-processed ROIs using MATLAB’s `detectMSERFeatures` function. The Maximally Stable Extremal Regions (MSER) algorithm detects stable intensity regions across multiple threshold levels, making it well suited for spheroid detection with their intensity variations caused by cells at different focal depths. Cell and spheroid segmentation was further refined by using the activecontour iterative image segmentation function in MATLAB. Contour points of the segmented spheroids and cells were determined with the bwboundaries MATLAB function. Shape parameters of the segemented cells and spheroids were then computed using regionprops. The spheroid speed in each frames was estimated by a central difference approximation. The spheroid speed in the inlet channel *V_in_*and outlet channel *V_out_*were calculated by averaging the approximately-constant spheroid velocity in the inlet channel (before entering the extension flow) and the outlet channel (after exiting the extension flow), respectively. The initial spheroid diameters *D*_0_ were the average equivalent diameter 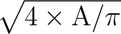 of the 2D projected area *A* in all frames in the inlet channel prior to the spheroid entering the extensional flow. Because spheroids were irregular in shape, tracking how their area was distributed in space via area moments of inertia was an appropriate approach to measure spheroid deformation [30]. The inertia ratio used to assess spheroid deformation was 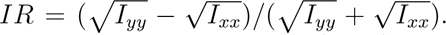 In each frame, the spheroid area moments of inertia were computed from all contour points (*x_i_, y_i_*) along with the shape centroid (*x_COM_, y_COM_*): *I_xx_* = ∑(*x_i_ − x_COM_*)^2^ and *I_yy_* = ∑(*y_i_ − y_COM_*)^2^. In that case, the closeness changes the force they are experiencing in the cross-slot so they were discarded. Only movies with one in-focus spheroid at a time inside the cross-slot were considered. Movies were discarded from analysis if spheroids were within one spheroid diameter of each other, which affected the hydrodynamic forces felt by the spheroids, spheroid were out-of-focus, and if spheroid rigidly rotated wihtout much stretching in the cross-slot.

### DIC Analysis of Intra-spheroid Strain

Only spheroids undergoing pure deformation without noticeable rotation were analyzed with digital image correlation (DIC) to estimate strains. NCorr [41] was implemented within a for-loop to compute the displacement of subset centers across the spheroid between consecutive frames. For each frame, a region of interest (ROI) centered on the spheroid was defined and used for DIC analysis. The quasi-spherical spheroid shape selection criterion was important because DIC introduces larger errors near object boundaries, as strain calculation at each grid point depends on the displacement data of neighboring points. Points located at the edges have fewer neighboring data points available, reducing the accuracy of strain estimation. These errors become more pronounced when the spheroid boundary contains irregular or non-smooth features. Over the time intervals in which spheroids experienced extensional flow, cumulative displacements were computed by adding together the displacements between consecutive frames while subtracting rigid motion, ensuring that only displacement due to deformation was retained. Rigid motion was computed by fitting each frame’s displacement field to a rigid motion model consisting of translation and small rotation. The measured displacement field was modeled as *u* = *u_RBM_*+ *u_def_*, where *u_RBM_* is the rigid body motion and *u_def_*is the displacement due to deformation. For small rotations, the rigid body motion consisted of a rigid translation and a rigid in-plane rotation: *u_RBM_* = *t_x_ − ωy, v_RBM_* = *t_y_*+ *ωx*, where *t_x_* and *t_y_* are the rigid translations and *ω* is the in-plane rotation angle [80]. These three parameters *t_x_*, *t_y_*, and *ω* were computed by a least-squares fit to the displacements of all subset centers (*x_i_, y_i_*). For each frame, the fitted rigid body motion *u_RBM_* and *v_RBM_* were then subtracted from the NCorr measured displacements *u* and *v*, leaving only the displacement due to deformation, *u_def_* and *v_def_*. Because rigid-body motion does not produce strain, this rigid body motion step does not affect the estimated displacement gradients and strains [81].

The displacement gradients *∂u/∂x*, *∂u/∂y*, *∂v/∂x*, and *∂v/∂y* of the cumulative displacement field were estimated at each point using a local least squares fitting approach over small strain calculation windows in which displacement distributions can be approximated as a linear plane [82]. Green–Lagrangian strains *E_xx_*, *E_xy_*, and *E_yy_* were estimated from the displacement gradients:

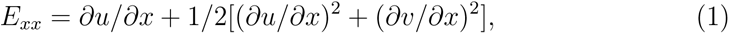

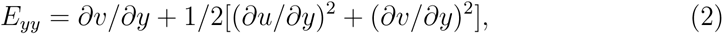

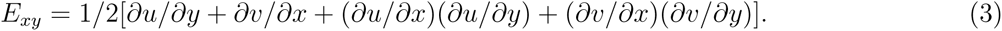

An equivalent 2D Von Mises strain was subsequently calculated from these strain components:

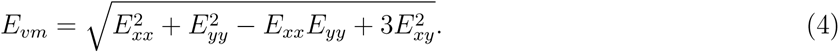

To compare deformation trends throughout the spheroid, Green-Lagrangian strains were compared across concentric annular regions. Large and small spheroids were divided into five radial layers based on the normalized distance from the spheroid centroid. The mean and standard deviation of the strain components (*E_xx_*, *E_yy_*, *E_xy_*) and equivalent strain (*E_vm_*) for each radial layer were computed from the DIC analysis subregions within that layer. Because DIC computes strain by plane fitting over a neighborhood of subset points, and subset centers near the spheroid boundary lack a full set of neighboring points, outer-layer strain estimates carry higher uncertainty; outer-layer strain was therefore computed only at positions with at least three neighboring subset centers available for plane fitting.

To compute deformation between the interior and periphery of each spheroid, an inner region was defined as all DIC subset centers with normalized radial distance of *r/D*_0_ ≤ 0.25 from the spheroid center, and an outer region as those with normalized distances *r/D*_0_ ≥ 0.6. These thresholds were chosen to reflect the inner and outer region deformation trends of the spheroid while excluding the intermediate region, while also ensuring a sufficient number of subset centers in each region. As a homogeneous, isotropic control for this inner/outer deformation comparison, 0.5% w/w agarose hydrogel particles were subjected to cross-slot extensional flow and DIC analysis as the spheroids. Hydrogel particles (diameter 25–50 µm) were deformed in a cross-slot device with a channel width of 290 µm and a depth of 265 µm. The characteristic extensional strain rates for the hydrogel microparticles ranged from Ω = 234.6 to 342.9 *s^−^*^1^, corresponding to a characteristic extensional force (*µ*Ω) of 25.8–37.7 Pa, where *µ* = 110 mPa·s is the viscosity of the buffer fluid.

### Live Spheroid Staining and Confocal Imaging

To measure spheroid morphology, live cell imaging was performed after 4 days of culturing for spheroids minimally adhered to the glass substrate, thereby mimicking their suspended state in microfluidic biomechanics experiments. Spheroid pellets were resuspended in warm media and deposited in a clear-bottom 96-well plate (Fisherbrand, Fisher Scientific). Following 30 minutes of settling time, spheroids were incubated for 50 minutes in a live stain solution consisting of a plasma membrane dye (CellMask Plasma Membrane Stain Orange, Invitrogen), a F-actin dye (CellMask Actin Tracking Stain Green, Invitrogen) and a DAPI stain (NucBlue Live ReadyProbes Reagent Hoechst 33342, Invitrogen) added to cold media. After a PBS wash, spheroids wells were replenished with DPBS containing calcium for imaging. Spheroid images and *z*-stacks were acquired using a Zeiss LSM 880 confocal laser scanning microscope (20× air-immersion objective). During imaging, spheroids were maintained in an enclosed incubation chamber at 37 *^◦^*C and 5% CO_2_. The pinhole was set to 1 Airy Unit. Imaging was performed using 405 nm (Diode), 488 nm (Argon), and 561 nm (DPSS) laser lines with relative laser powers of 0.03, 0.024, and 0.02, respectively. Single images were acquired at the spheroid midplane in 8-bit mode with frame averaging of 4 and a frame size of 1024 × 1024 pixels. *Z*-stack images encompassing the entire spheroid volume were acquired with a *z*-step (voxel depth) of 0.7348177 µm, frame averaging of 1, and a frame size of 512 × 512 pixels.

### Cell and Nuclear Morphology Analysis

Individual cell morphology within spheroids was assessed in two ways. First, single 2D midplane images of small and large spheroids were analyzed for cell area and circularity based on plasma membrane staining. Second, 3D spheroid *z*-stacks were analyzed using three-dimensional segmentation to identify individual nuclei and quantify nuclear volume and sphericity.

Cell area and circularity were quantified from plasma membrane confocal images. For small spheroids, cell boundaries of all cells were manually outlined using the freehand tool in Fiji across multiple *z*-slices in the the slice corresponding to each cell’s center plane. For large spheroids, the high cell density made analysis of all cells impractical. Therefore, the mid-plane of the *z*-stack was selected, and in-focus cells within that plane were manually segmented. Segmented cell region circularity and area were measured from the Fiji (ImageJ 1.54p; [83]) Measure tool. For comparison, isolated single cells suspended in a viscous buffer were imaged by brightfield microscopy (10× and 40× objectives, Prime95B camera), and cell area and circularity were quantified from binarized single-cell masks in MATLAB using the regionprops function.

Nuclei volumes were measured as a proxy for cell volumes, given that the cell-to-nuclear volume ratio is approximately constant [43, 21, 48]. Individual nuclei within spheroids were identified from confocal *z*-stacks using Cellpose 3D segmentation (v2.2.2,`"nuclei"` pretrained model; `channels=[0,0]`; `do_3D=True`; anisotropy set to the ratio of the *z*-step to the *xy* pixel size for each stack) [84] in a custom three-dimensional image analysis workflow implemented in Python (Google Colab), after smoothing and intensity normalization of the *z*-stacks. For each segmented nucleus, the 3D centroid (center of mass of the voxel intensity distribution) and volume (voxel count converted to cubic micrometers) were computed, and nuclear sphericity *psi* was calculated in MATLAB from the segmented nucleus volume *V* and surface area *A*: *ψ* = [*π*^(1^*^/^*^3)^(6*V*)^(2^*^/^*^3)^]*/A*. Segmented objects with a volume < 100 µm^3^ (equivalent diameter ∼ 6 µm), or spanning fewer *z*-slices than required for a 6 µm-diameter object, were excluded as noise. Segmentation quality was assessed by visual inspection of all detected nuclei, with *>* 90% correctly segmented. The spheroid 3D centroid was defined as the mean position of all nuclear centroids, and each nucleus’s normalized distance was calculated relative to this centroid. For the core-shell nuclear volume comparison, the core was defined as nuclei with normalized distance *r/D*_0_ < 0.55 and the shell as nuclei with normalized distance *r/D*_0_ *>* 0.7; This inner threshold was expanded relative to the DIC strain analysis to ensure a sufficient number of nuclear per region.

Live cell confocal images were optimized for display in figures using the ImageJ Hi-Lo lookup table. All analysis was performed on unedited images.

### Immunofluorescence Staining and Confocal Imaging

To assess protein distributions within spheroids, day-4 spheroids were fixed and stained for vimentin, E-cadherin, nuclei, and F-actin. Immunofluorescence staining was performed on intact 3D spheroids without embedding, preserving their three-dimensional structure. Spheroids were fixed with buffered formalin (diluted 1:10), permeabilized with 0.5% (v/v) Triton X-100 in PBS, and quenched with 0.1 M glycine. Spheroids for vimentin and E-cadherin immunostaining were then blocked with StartingBlock (PBS) Blocking Buffer (Thermo Fisher Scientific, 37538) before antibody incubation. In a clear-bottom 96-well plate (Fisherbrand, Fisher Scientific), spheroids were incubated overnight at 4 *^◦^*C with primary antibodies (diluted 1:200) against E-cadherin (4A2 Mouse Monoclonal Antibody, #14472, Cell Signaling Technology) and vimentin (D21H3 Rabbit Monoclonal Antibody, #5741, Cell Signaling Technology), or with ActinGreen™ 488 ReadyProbes™ Reagent (Invitrogen, Thermo Fisher Scientific, R37110) for F-actin staining. After PBS washing the following day, antibody-stained spheroids were incubated for 2 hours at room temperature with secondary antibodies diluted 1:1000 (anti-rabbit Alexa Fluor 488, Cell Signaling Technology; anti-mouse Alexa Fluor 568). Spheroids were mounted with ProLong™ Diamond Antifade Mountant with DAPI (Thermo Fisher Scientific) for nuclear staining and stored at 4 *^◦^*C overnight prior to imaging. Confocal imaging was performed using a Zeiss LSM 880 confocal microscope with 20× air and 40× oil-immersion objectives and a pinhole set to 1 Airy unit. Spheroids were imaged using three laser lines: 405 nm (diode), 488 nm (argon), and 561 nm (DPSS). The relative laser power settings in the microscope software were 0.03, 0.022, and 0.02, respectively. Laser power, detector gain, and pinhole settings were kept constant for all samples. Images were acquired in 8-bit mode. For each spheroid, a single slice at mid-depth and a *z*-stack spanning the entire spheroid (voxel depth 0.73 µm) were acquired, using line averaging of 8 for single-slice images and 1 for *z*-stacks. The specificity of the primary antibodies against vimentin and E-cadherin was verified with appropriate control conditions.

### Immunofluorescent Image Analysis

Protein level analyses were preformed on approximately one-cell-thick (10 µm) sum-slice *z*-projections around the spheroid mid-depth location. Spheroid equivalent diameters *D*_0_ and centroids were calculated from the binarized 2D projections of the whole spheroid, three-channel *z*-stack. Multi-channel confocal *z*-stack tiff files were imported into MATLAB using the Bio-Formats toolbox [85]. Nucleus, vimentin, and E-cadherin channel images were separately background-subtracted, smoothed (standard deviation = 1), normalized to [0 1], and binarized using Otsu’s method using MATLAB functions. Spheroid mid-depth region 2D projections were divided into concentric layers approximately 10 µm in thickness from outer spheroid edge inward. (The spheroid outer edge location was estimated from the segmented spheroid shape from the region’s sum-slice *z*-projection of all three channels.) For each layer, the average and standard deviation of intensity across all positive pixels were computed for nuclei, vimentin, and E-cadherin. F-actin distribution was visually assessed. Immunofluorescent images were adjusted for brightness and contrast using ImageJ HiLo lookup tables for visualization purposes only. All quantitative analyses started from the original, unprocessed images.

### Statistical Analysis

Pairwise comparisons between two independent groups (e.g., small vs. large spheroids) were performed using the Mann-Whitney U (Wilcoxon rank-sum) test. Paired comparisons between two regions of the same spheroid (e.g., inner vs. outer layers) were performed using the Wilcoxon signed-rank test. Comparisons across three or more groups were first evaluated with the Kruskal-Wallis test, followed by pairwise post-hoc comparisons using Dunn’s test with Bonferroni correction for multiple comparisons. These non-parametric tests were used throughout because of small sample sizes, which limited reliable assessment of normality. A significant threshold of *p <* 0.05 was used throughout, and all statistical tests were performed in MATLAB. All violin distribution plots were generated using the Violinplot-Matlab toolbox [86].

## Supporting Information

Supporting Information is available from the Wiley Online Library.

## Acknowledgments

This study was supported by the National Science Foundation (CMMI Award No. 2301804), the College of Science and Mathematics Dean’s Doctoral Research Fellowship (funded by Oracle, Project ID R20000000025727), and the Henry Luce Foundation (Project ID S20240000058705). We thank the UMass Boston Bioimaging Core Facility for support with confocal imaging, and Dr. Shakir Khan (UMass Boston and Mass General Hospital) for guidance in cell culture and handling.

## Conflicts of Interest

The authors declare no conflicts of interest.

## Notes

### Competing Interest Statement

The authors have declared no competing interest.

