## Supplementary Information for "Extensional Flow Microfluidics Reveal Size-Dependent Core-Shell Mechanical Heterogeneity in Pancreatic Tumor Spheroids"

### 1 Supplementary Methods

#### 1.1 Viscous buffer viscosity measurement

The same viscous buffer was used in all deformation experiments, consisting of 0.8% w/v methylcellulose (Thermo Scientific Chemicals, CAS 9004-67-5, molecular weight 454.513 g/mol) dissolved in DPBS (1×) containing calcium (Gibco). The buffer was filtered by vacuum filtration using grade 5 cellulose paper filter with a pore size of 2.5 textmu m.

A TA Instruments Discovery Hybrid series rheometer was used to measure the buffer viscosity. Shear rate sweep tests were conducted using a 40 mm parallel plate geometry over a shear rate range of 1–1000 $s^{-1}$ with 10 points per decade. The shear rate sweep test was repeated three times.

The viscous buffer exhibited shear thinning behavior, meaning viscosity decreases as shear rate increases. To determine the effective viscosity relevant to the microfluidic deformation experiments, the zero-shear viscosity was estimated from the average of the three rheology replicates. As discussed in our previous work [1], the dominant shear rates in the cross-slot microfluidic device, represented by the off-diagonal velocity gradients ($\partial u/\partial y$ and $\partial v/\partial x$), are approximately zero near the stagnation point. Therefore, viscosity measurements at low shear rates most representative of the fluid conditions within the cross-slot.

However, for parallel plate geometries, the first reliable rheological measurements are typically obtained as low as shear rates of approximately 10 $s^{-1}$. Buyukurganc et al. [2] demonstrated that viscosity measurements at shear rate of 10 $s^{-1}$ agree well with zero-shear viscosity measurements obtained using concentric cylinder geometries, which provide more accurate measurements at low shear rates for methylcellulose solutions.

Accordingly, the buffer viscosity was calculated by averaging the measured viscosities at shear rates of 10 and 12.6 $s^{-1}$ across the three rheology replicates, resulting in an estimated zero-shear viscosity of 90 mPa·s.

#### 1.2 Digital Image Correlation Validation

NCorr DIC analysis for measuring strain across spheroids was validated by applying known digital deformations to spheroid images and comparing the NCorr DIC measurements with analytical expectations.

##### 1.2.1 10% Strain

Because the Von Mises strain is primarily influenced by the strain along the $y$-axis ($E_{yy}$), which is the direction of the outlet channel and the principal stretching direction in extensional flow, we applied known deformations along the $y$-axis to an undeformed spheroid and computed the resulting strain using NCorr DIC (**Figure S8**). **Figure S8a** shows a large spheroid in its undeformed state and after applying 10% strain about the spheroid center along the $y$-axis using an affine transformation in MATLAB. The cumulative displacement resulting from this imposed deformation was computed using NCorr DIC (**Figure S8b**), and the corresponding strain components are shown in **Figure S8c**.

As observed in the strain maps, $E_{xx}$ and $E_{xy}$ are approximately zero (on the order of $10^{-4}$), as expected, while $E_{yy}$ has values near 0.105, consistent with $E_{yy}$ expected with this deformation:

$$x' = x, \quad y' = 1.1y \tag{1}$$

$$u(x,y) = 0, \quad v(x,y) = 0.1y \tag{2}$$

$$\frac{du}{dx} = 0, \quad \frac{du}{dy} = 0 \tag{3}$$

$$\frac{dv}{dx} = 0, \quad \frac{dv}{dy} = 0.1 \tag{4}$$

$$E_{yy} = \frac{dv}{dy} + 0.5\left(\left(\frac{du}{dy}\right)^2 + \left(\frac{dv}{dy}\right)^2\right) \tag{5}$$

$$E_{yy} = 0.1 + 0.5(0.01) = 0.105 \tag{6}$$

**Figure S8d** shows the key strain components in this case ($E_{yy}$ and $E_{vm}$) across radial layers of the spheroid, demonstrating that they remain nearly constant at approximately 0.105 across all layers.

We deformed the spheroid along the $y$-axis using four prescribed strain levels and compared the NCorr DIC measurements of the average $E_{YY}$ with the applied deformations (**Figure S8e**). The data points follow a linear trend, with a fitted slope of 1.061 and an $R^2$ value close to 1.

##### 1.2.2 Rigid Rotation and Translation

To further validate the NCorr DIC results, we evaluated strain measurements under rigid body motion of the spheroid, including small rotations, and pure translations. In both cases, no deformation is expected; therefore, all strain components should be approximately zero.

The NCorr DIC results for rotating the spheroid about its center by a small angle of 2° are shown in **Figure S9a and S9b**. All strain parameters are approximately zero, as expected. The average *pm* standard deviation of strains across radial layers of the spheroid (**Figure S9c**) also confirms near-zero strain values. The average $E_{vm}$ from four small rotations is plotted in **Figure S9d**, showing a linear fit with a slope of zero, indicating negligible strain across all cases.

For validating translation motions, the spheroid was translated using four prescribed equal displacements with $dx = dy$. **Figure S10a** shows the cumulative displacement for the case of $dx = dy = 10$ pixels (i.e., the spheroid shifted 10 pixels to the right and 10 pixels downward). The average *pm* standard deviation of strain across radial layers of the spheroid is shown in **Figure S10b**, where all values are approximately zero, as expected. Finally, a linear fit was performed between the average $E_{vm}$ computed by NCorr and the four applied translations ($dx = dy$). The resulting fit has a slope of zero (Figure S10c), confirming negligible strain under pure translation.

##### 1.2.3 Shear Strain

We further validated our DIC results for shear strain by applying four known shear strains (0.02, 0.05, 0.1, and 0.2) in the $y$-direction to the undeformed spheroid (**Figure S11**). In the microfluidic experiments, the spheroid undergoes a small shear deformation when transitioning from inlet-directed motion to outlet-directed motion in the cross-slot. Since the outlet channel is aligned with the $y$-axis, the shear strains were applied in the $y$-direction. This validation assesses the sensitivity of DIC in detecting spheroid shear deformation. The shear strains were applied using a 2D affine transformation in MATLAB. Based on the Green-Lagrangian strain definition, we expected shear strain component ($E_{xy}$) is equal to one half of the applied known shear strain.

**Figure S11a** shows the displacement arrows for a spheroid subjected to a known shear strain of 0.1 (10%) in the $y$-direction. **Figure S11b** presents the DIC strain measurements across radial layers of the spheroid for this known shear deformation. For a known shear strain of 0.1, the expected Green-Lagrangian strain components are $E_{xx} = 0.005$, $E_{yy} = 0$, $E_{xy} = -0.05$, and $E_{vm} = 0.0867$. The measured strains in **Figure S11b** closely match these expected values, with only minimal variation from the mean strains (all standard deviations were on the order of $10^{-4}$. The shear strain ($E_{xy}$) is negative because the right side of the spheroid moves toward the negative $y$-direction. In the image coordinate system, the $y$-axis is reversed; therefore, motion in the negative $y$-direction corresponds to arrows pointing upward.

Based on the Green-Lagrangian strain equation for $E_{xy}$, the expected value is equal to one-half of the applied shear strain. We plotted the NCorr DIC measurements of $E_{xy}$ against the known applied shears strains and fitted a linear regression model (**Figure S11c**). The data showed an excellent linear relationship, with a slope of approximately 0.5 and an $R^2$ of 1.

### 2 Supplementary Tables

A summary of the experimental conditions for all groups is provided below:

Table S1: Experimental parameters for each group.

| | **Single cells** | **Small spheroids** | **Large spheroids** |
|---|---|---|---|
| Mean $D_0$ (µm) | 20.17 | 55.18 | 135.13 |
| Range of $D_0$ (µm) | 14.24–25.51 | 38.61–73.85 | 99.73–158.68 |
| $N$ number | 19 | 22 | 27 |
| Channel width $w$ (µm) | 70 | 150 | 264 |
| Channel depth (µm) | 55 | 137 | 268 |
| Flow rate $Q$ | 324 µL/hr | 4.3 mL/hr | 30 mL/hr |
| Flow ratio (sheath:center) | 1 | 1 | 1 |
| Frame rate (fps) | 2200 | 3000 | 3200 |
| Exposure time (µs) | 2.126 | 1.6 | 1.885 |

### 3 Supplementary Movies

**Movie S1.** Time-lapse movie of the representative PANC-1 single cell stretching experiment. Time zero is when the single cell enters the extensional flow region (velocity starts decreasing from its steady-state inlet velocity value).
**Movie S2.** Time-lapse movie of the representative PANC-1 small spheroid stretching experiment. Time zero is when the spheroid enters the extensional flow region.
**Movie S3.** Time-lapse movie of the representative PANC-1 large spheroid stretching experiment. Time zero is when the spheroid enters the extensional flow region.
**Movie S4.** Time-lapse movie of a large spheroid showing several cells attached to one side of the spheroid become significantly stretched outside the spheroid bulk (time zero = entry into the extensional flow region).
**Movie S5.** Time-lapse movie of a spheroid showing rigid rotation in the cross-slot and no stretching (time zero = entry into the extensional flow region).
**Movie S6.** Time-lapse movie of a large spheroid showing substantial stretching and separation of several cells that were loosely attached to the spheroid (time zero = entry into the extensional flow region).
**Movie S7.** Time-lapse movie of a small spheroid showing the rupture of one or more cells in the outlet channel, indicated by a line extending from the spheroid containing the remnants of the dead cell(s) (time zero = entry into the extensional flow region).

### 4 Supplementary Figures

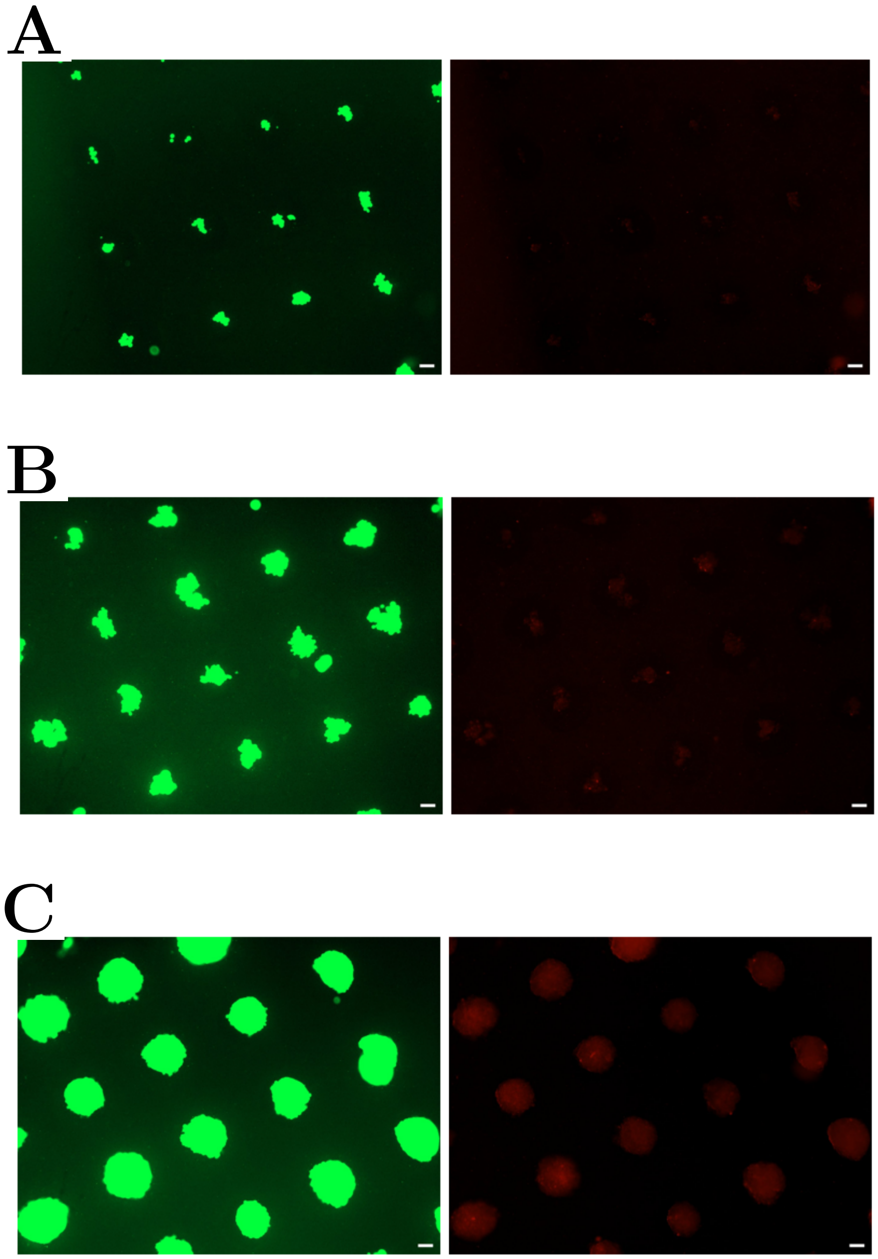


Figure S1: Fluorescence images of spheroids viability (green = live cells, red = dead cells) on **(A)** day 1, **(B)** day 4, and **(C)** day 12. These days were selected to show the trend. Scale bars represent 100 µm.

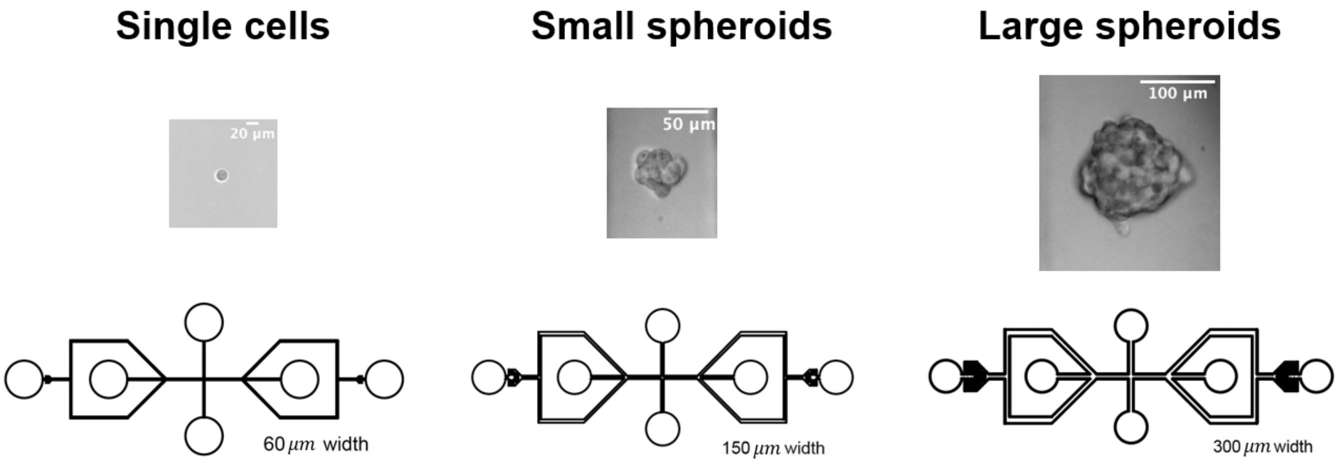


Figure S2: Spheroid and cell stretching was achieved by exploiting the extensional forces generated within cross-slot microfluidic devices. For each group, the intended square channel width was three times the targeted spheroid diameter (60, 150, and 300 µm widths, respectively). All devices incorporated sheath flow focusing.

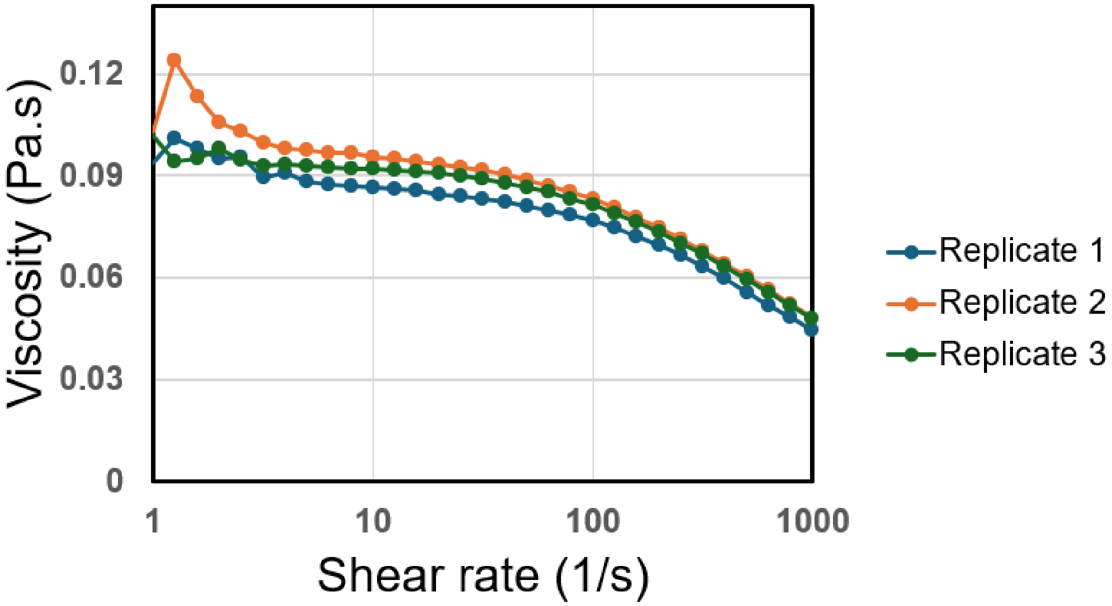


Figure S3: Viscosity versus shear rate for three replicate shear rate sweep measurements of 0.8% w/v methylcellulose dissolved in DPBS (1×) containing calcium. Measurements were performed using a TA Instruments Discovery Hybrid rheometer with a 40 mm parallel plate geometry.

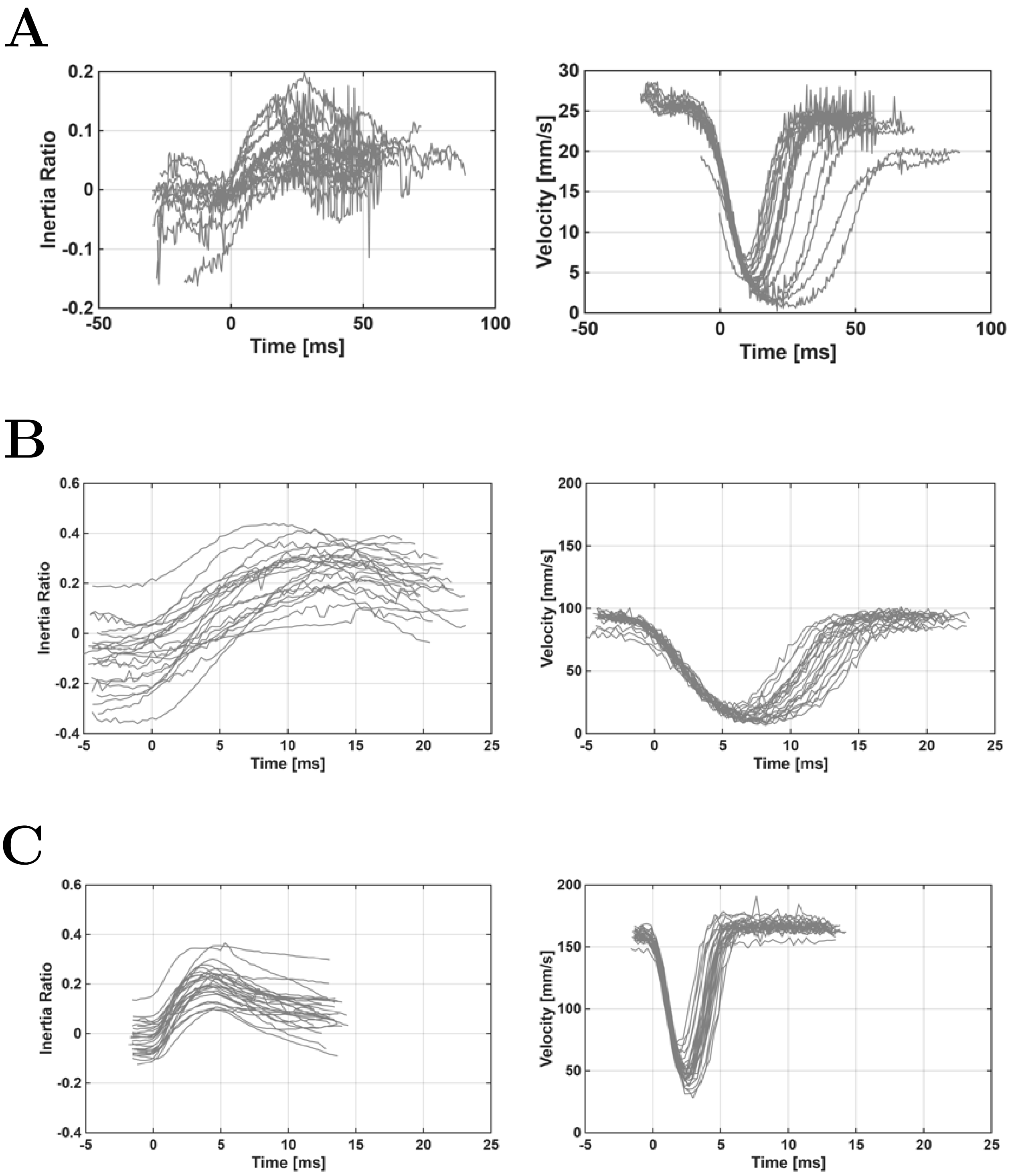


Figure S4: Shape change analysis from 2D contours of suspended spheroids and single cells stretching in microfluidic extensional flow. Inertia ratio and velocity versus time for all **(A)** single cells (mean diameter 20.17 µm; range: 14.24–25.51 µm; $N = 19$), **(B)** small spheroids (mean diameter 55.18 µm; range: 38.61–73.85 µm; $N = 22$), and **(C)** large spheroids (mean diameter 135.13 µm; range: 99.73–158.68 µm; $N = 27$) that underwent deformation without rotation or motion of loosely attached cellular components. Time zero corresponds to the moment the spheroid enters the extensional flow region. The plots includes the time spent in a portion of the inlet channel, the extensional flow region, and a portion of the outlet channel.

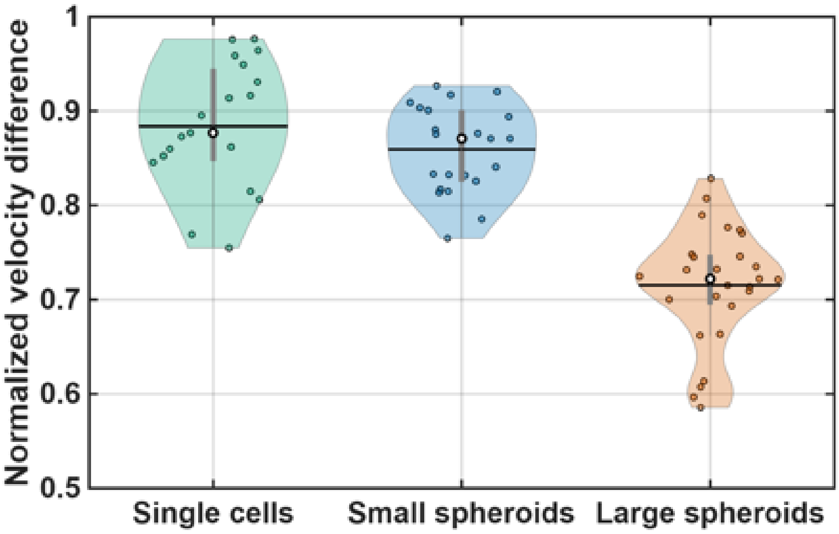


Figure S5: How closely the cells and spheroids approached the stagnation point was quantified from the normalized change in velocity $(V_{out} - V_{min})/V_{out}$ between the straight channel inlet speed and the minimum speed $V_{min}$ near the stagnation point. Normalized velocity difference showed that single cells and small spheroids slowed down more than large spheroids, though most objects were within $(V_{out} - V_{min})/V_{out} > 0.7$ and therefore felt similar hydrodynamic extensional forces [3].

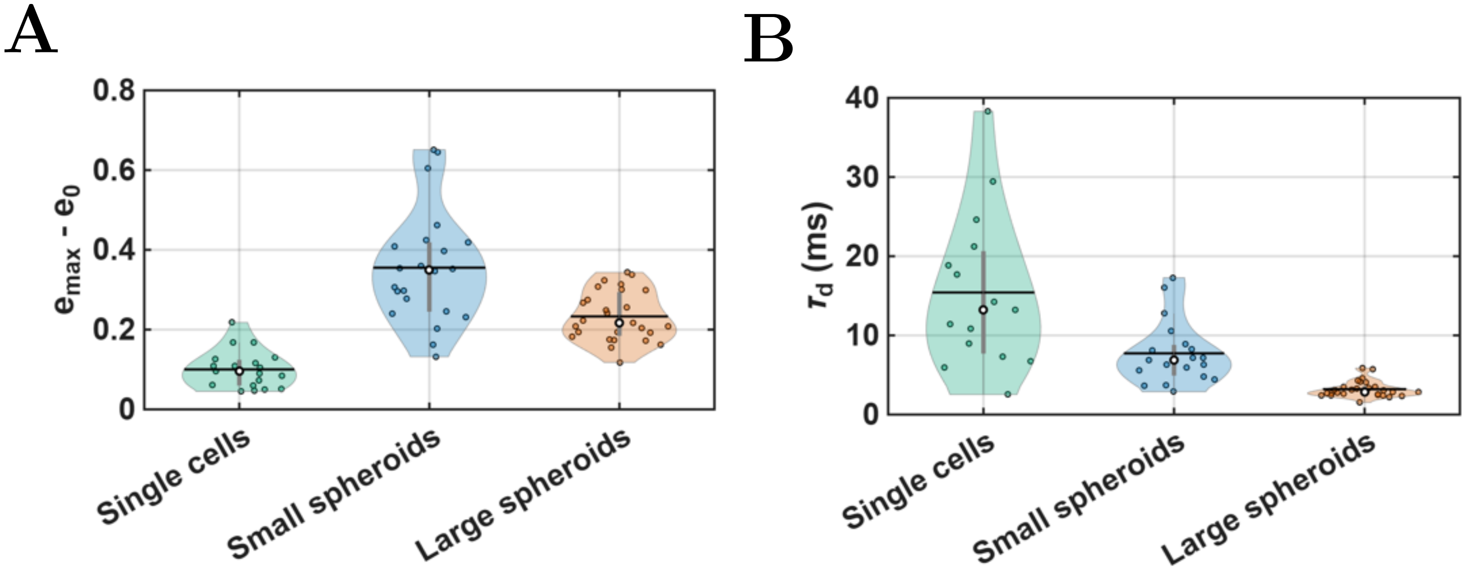


Figure S6: Shape change analysis from 2D contour shape for spheroids and single cell stretching in microfluidic extensional flow. **(A)** Difference in maximum inertia ratio and initial inertia ratio. Single cells stretched less compared to spheroids. The initial inertia ratio ($e_0$) was the average the inertia ratio in the inlet channel, and the maximum inertia ratio ($e_{max}$) was the peak value during extensional flow. **(B)** The deformation time constant ($\tau_d$) was computed by fitting an exponential function to the inertia ratio deformation data, starting from the onset of extensional flow and deformation to the point of maximum inertia ratio. The differences can be explained from the experimental conditions, where microfluidic device flow rates increased from single cells to small spheroids to large spheroids in order to maintain similar fluid extensional strain rates in devices of increasing size. Violin plots: The median is indicated by a white circle outlined in black, and mean is shows as the horizontal black line.

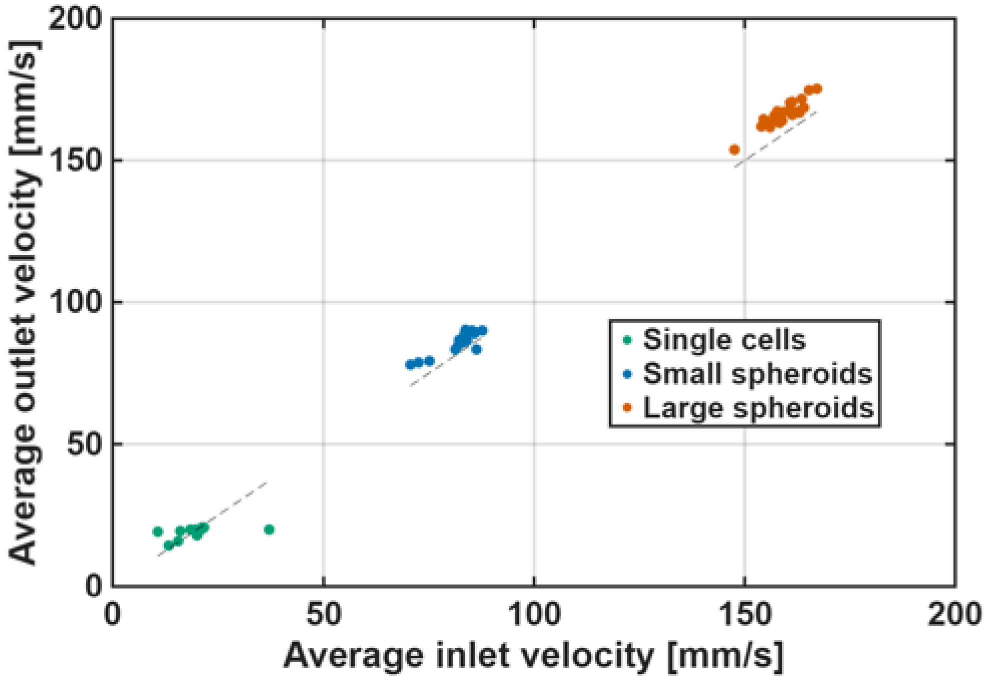


Figure S7: Comparison of steady-state average velocities in the inlet and outlet straight channels. Velocities were averaged in the inlet prior to entering the extensional flow and in the outlet after exiting the extensional region. The velocities in the inlet and outlet straight channels were nearly equal with data from all three experimental groups lying close to the dashed line $y = x$.

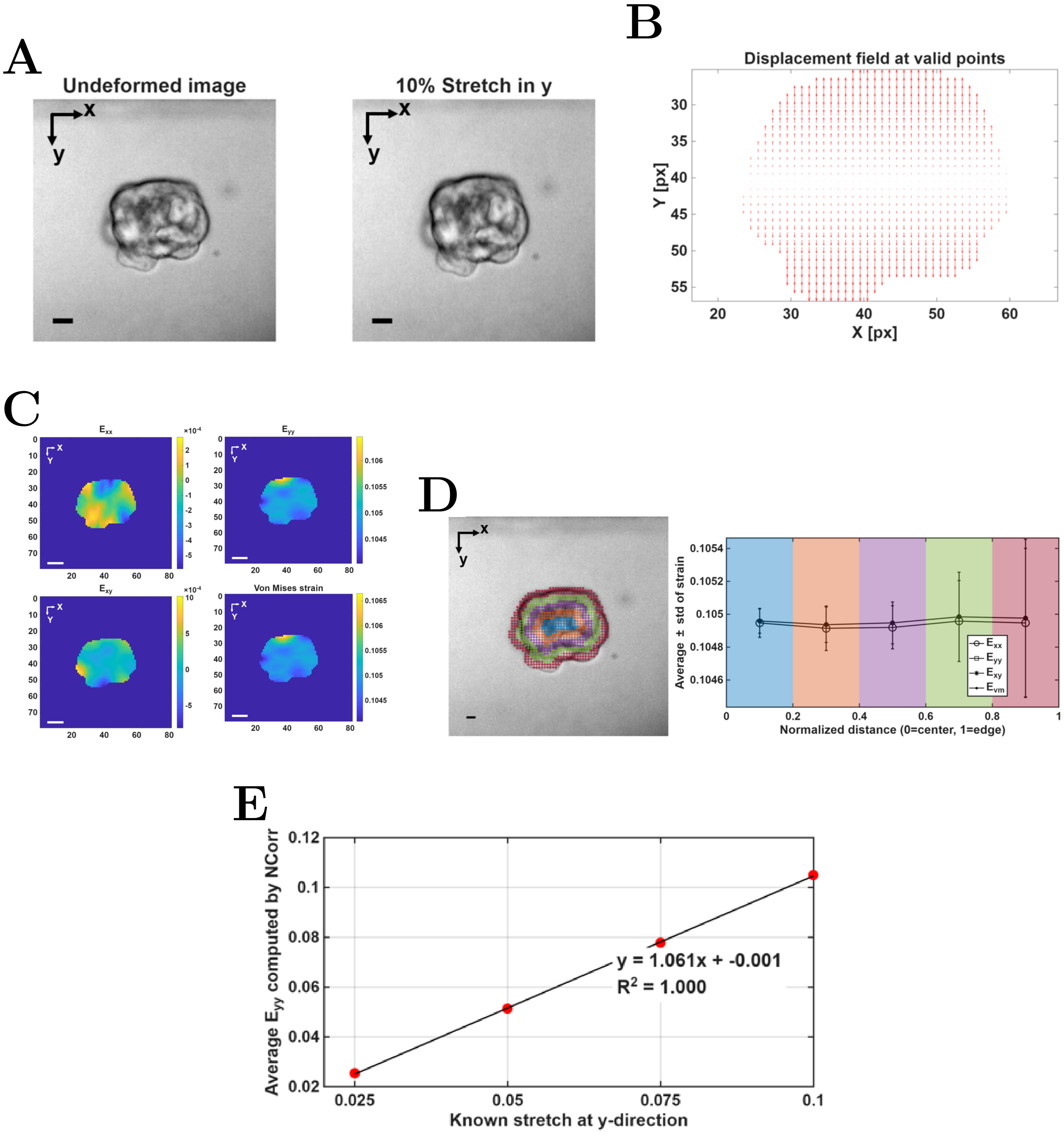


Figure S8: Validation results for NCorr DIC for a representative large spheroid only being stretched with 10% strain along the outlet ($y$-axis) for a known strain in the cross-slot. **(A)** Large spheroid before and after being stretched 10% at $y$-axis **(B)** Cumulative displacement arrows computed by NCorr DIC **C** Strain over x ($E_{xx}$), y ($E_{yy}$), shear strain ($E_{xy}$), and Von Mises strain ($E_{vm}$) computed for this known stretch. **D** Observing average ± standard deviation of $E_{yy}$ and Von Mises strain of various radial layers across the spheroid. **E** Comparison of the known stretch of spheroid over $y$-axis with the $E_{yy}$ computed by NCorr DIC for four known stretches. All scale bars equal 20 µm.

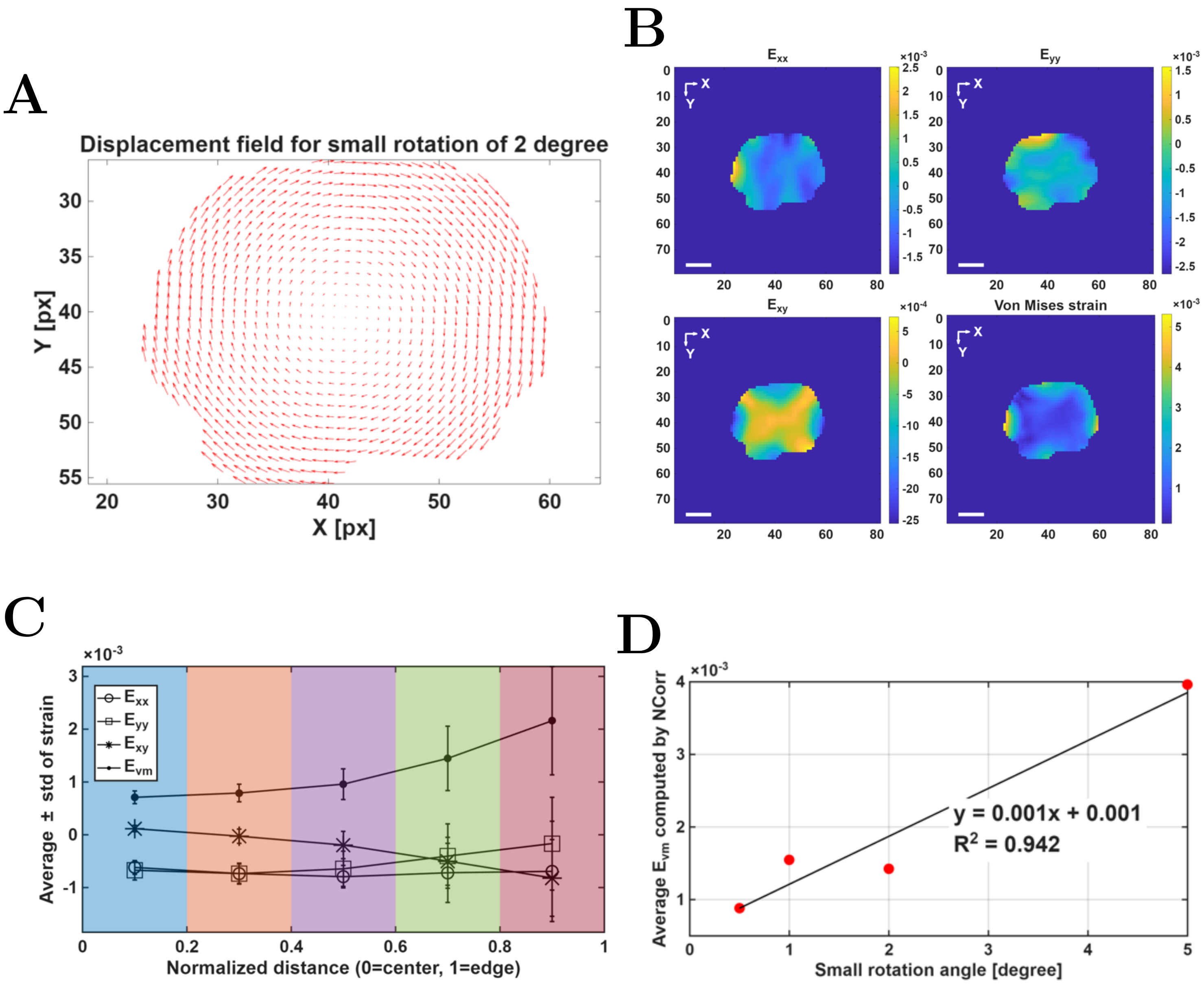


Figure S9: Validation results for NCorr DIC analysis for a representative large spheroid undergoing a 2° rigid rotation. **(A)** Cumulative displacement arrows computed by NCorr DIC for a small rotation of 2 degrees **(B)** Green-Lagrangian strain components in the $x$-axis ($E_{xx}$), $y$-axis ($E_{yy}$), and shear ($E_{xy}$) directions, and the von Mises equivalent strain ($E_{vm}$). **(C)** Observing average ± standard deviation of strains of various radial layers across the spheroid were on the order of $10^{-3}$, which was effectively zero and consistent with the imposed rigid rotation. **(D)** Comparison of the NCorr DIC average $E_{vm}$ for four known small rotations. All scale bars equal 20 µm.

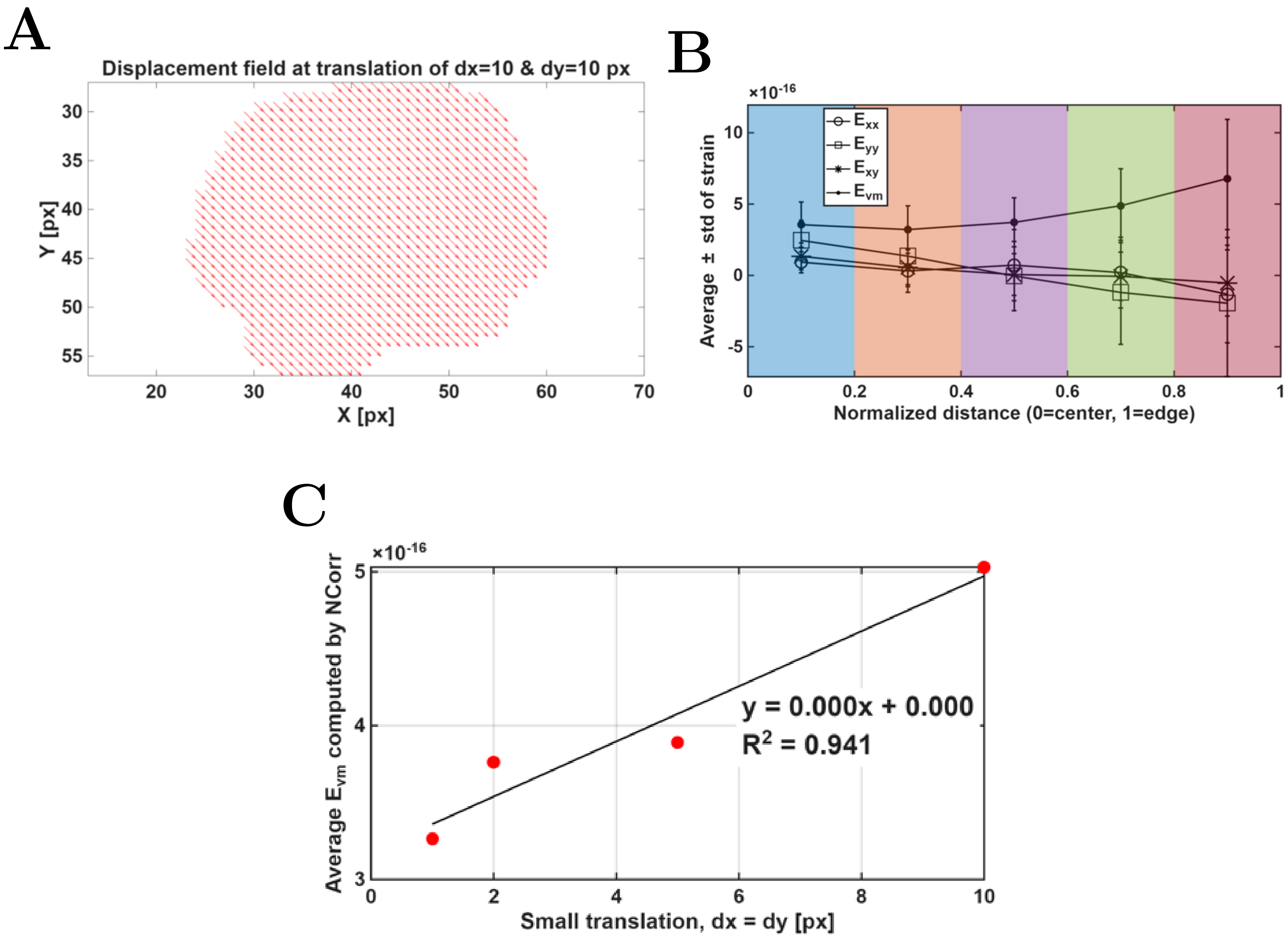


Figure S10: Validation results for NCorr DIC for a representative large spheroid undergoing pure translation ($dx = dy$). **(A)** Cumulative displacement arrows computed by NCorr DIC for translation of $dx = dy = 10$ pixels. **(B)** Observing average ± standard deviation of strains of various radial layers across the spheroid. The order of magnitude of these strains is $10^{-16}$, or practically zero, as expected for pure translation. **(C)** Comparison of the NCorr DIC average $E_{vm}$ for four known translations ($dx = dy$).

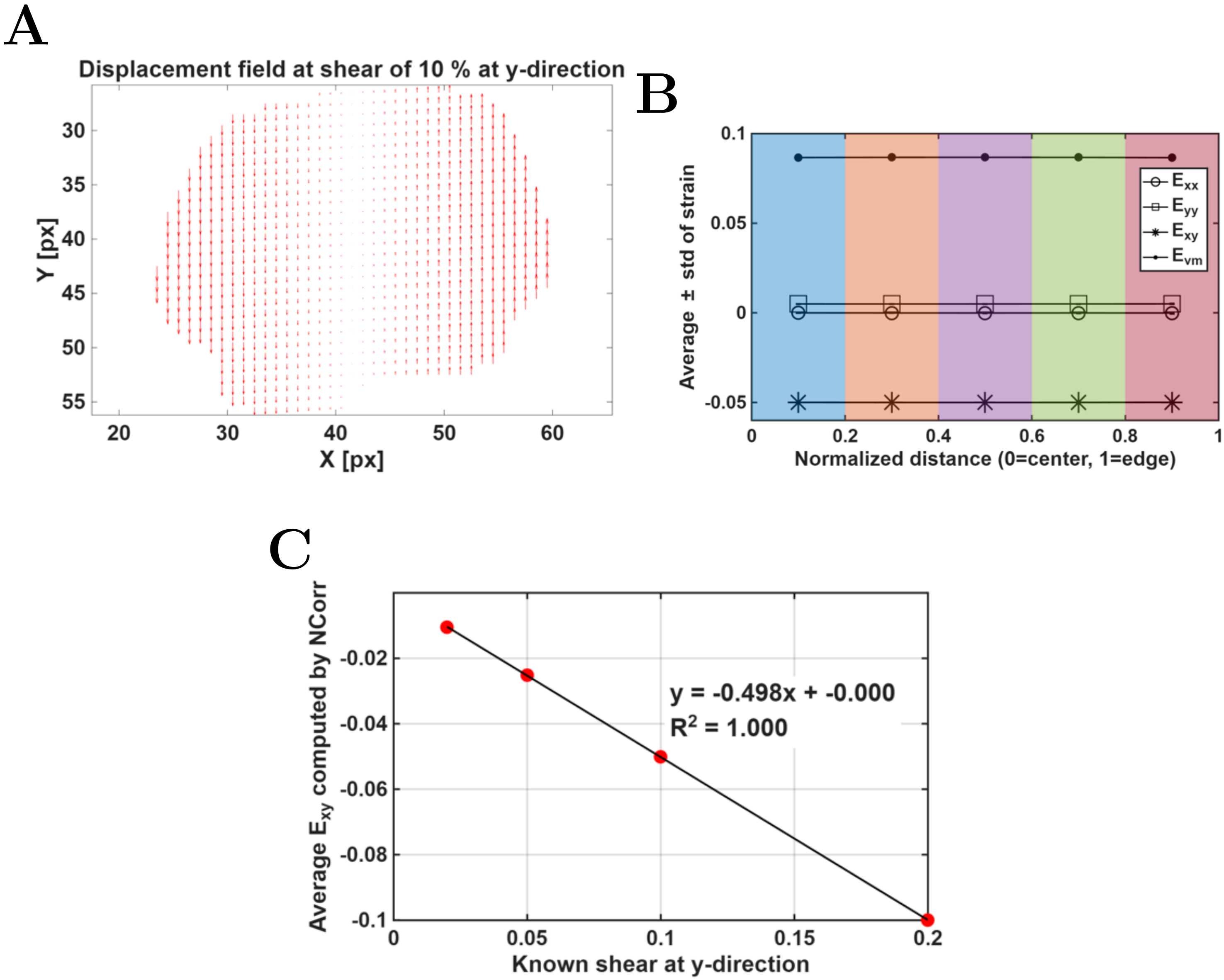


Figure S11: Validation results for NCorr DIC for a representative large spheroid undergoing shear at $y$-direction. **(A)** Cumulative displacement arrows computed by NCorr DIC for shear strain of 10 % at $y$-direction. **(B)** Observing average ± standard deviation of strains of various radial layers across the spheroid. **(C)** Comparison of the NCorr DIC average $E_{xy}$ for four known shears at $y$-direction.

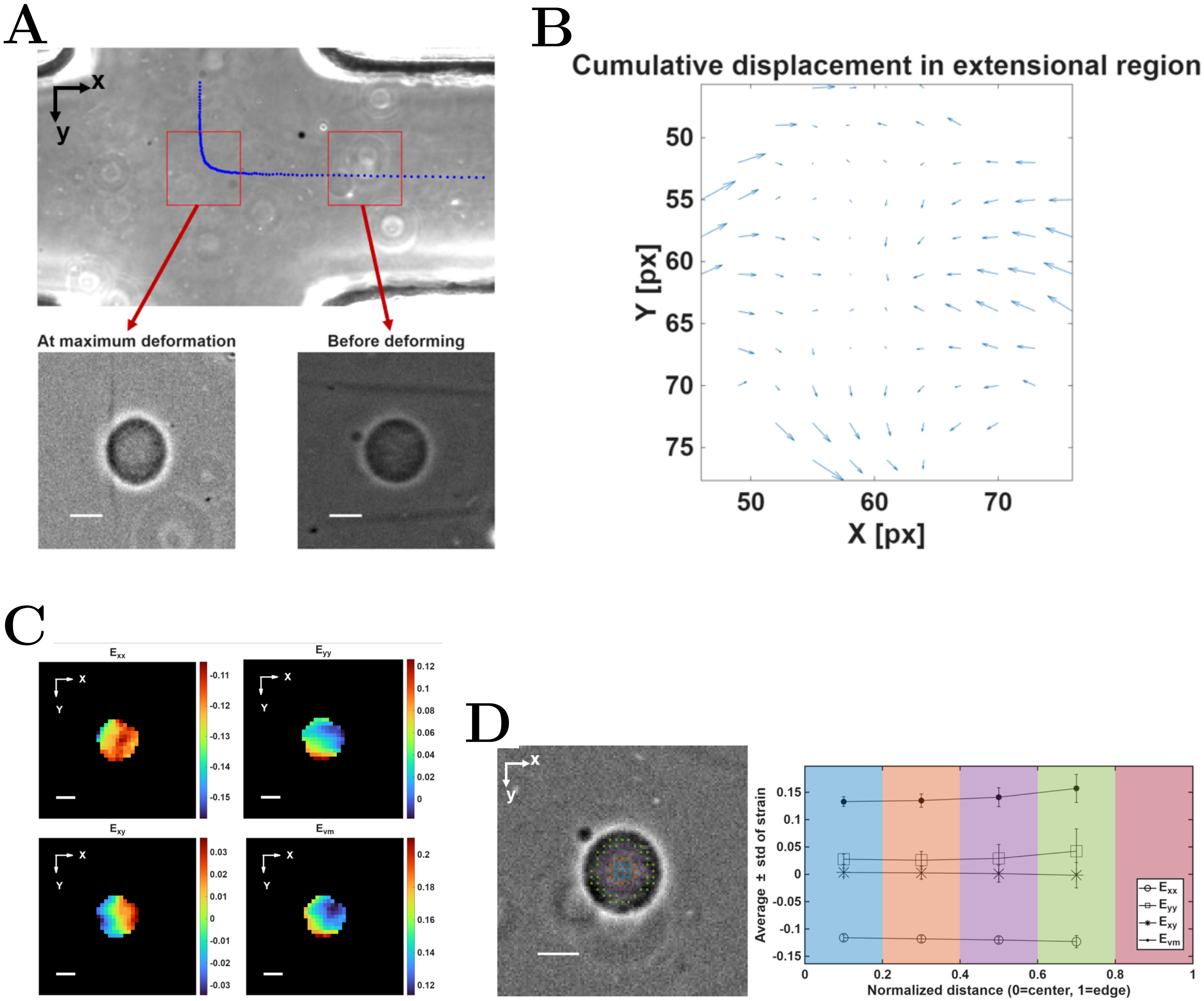


Figure S12: DIC analysis results for a representative hydrogel particle in the cross-slot. **(A)** Hydrogel particle at the onset of the extension region (immediately before deforming), and at its maximum deformation **(B)** Cumulative displacement arrows during deformation in the cross-slot **C** Strain over x ($E_{xx}$), y ($E_{yy}$), shear strain ($E_{xy}$), and Von Mises strain ($E_{vm}$). **D** Observing average ± standard deviation strain of various radial layers across the hydrogel. All scale bars equal 20 µm.

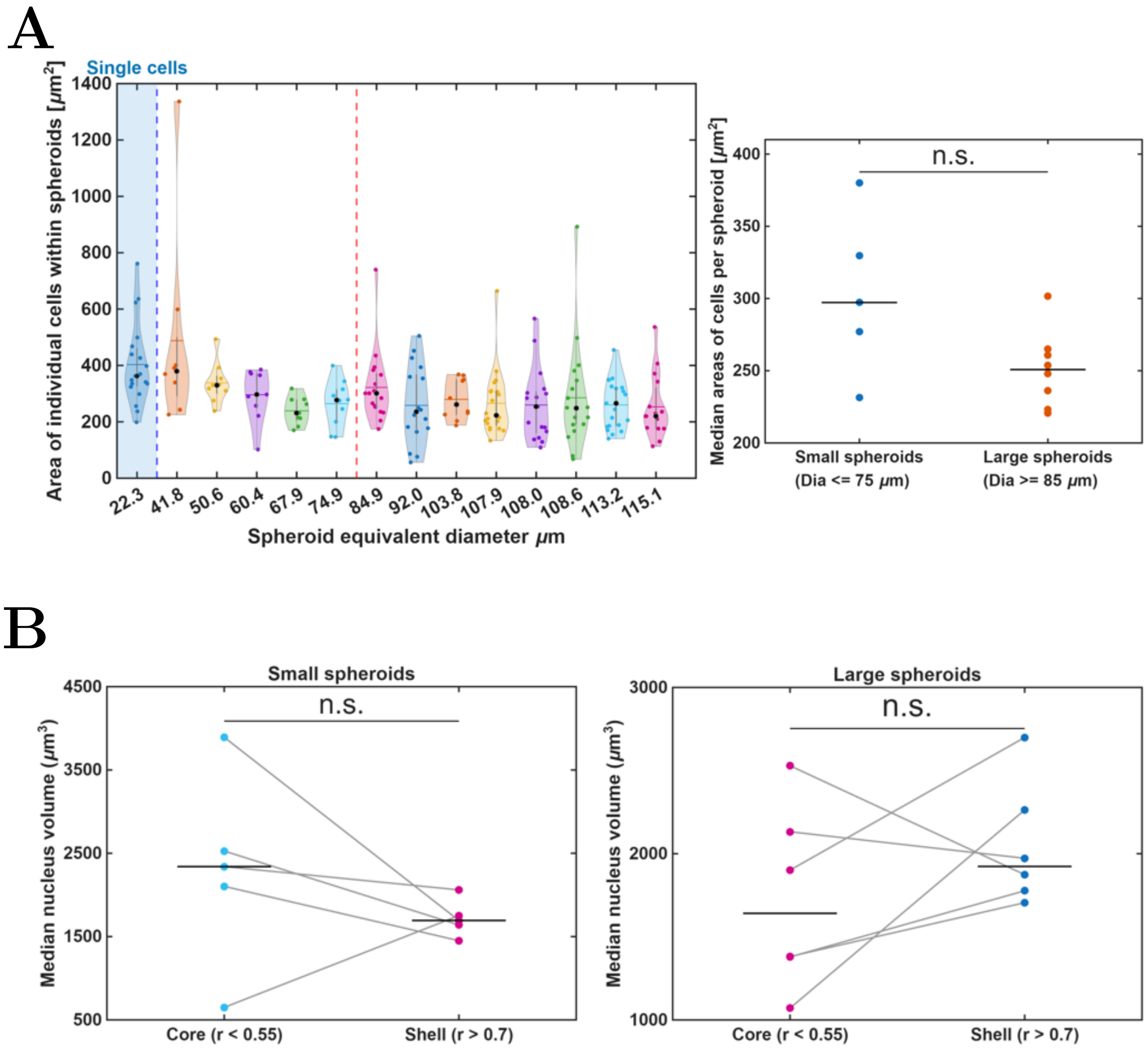


Figure S13: Morphology analysis of individual cells within small and large spheroids. **(A)** 2D cell area measurements. Blue shaded region: area of isolated single cells ($N = 19$) from brightfield images of suspended cells in a viscous medium. Unshaded region: Area of individual cells within 2D images of small with diameter 59.11 ± 13.22 µm (mean ± standard deviation, $N = 5$) and large spheroids with diameter 104.19 ± 10.48 µm ($N = 8$). Mann-Whitney U (rank-sum) test between median cell area per small and large spheroids, p = 0.093. **(B)** Median 3D nuclei volume for inner and outer regions (core and shell) of small (diameter 60.67 ± 13.84 µm, $N = 5$) and large spheroids (Diameter 103.45 ± 8.82 µm, $N = 6$). Core and shell is defined by the normalized nuclei distance ($r/D_0$) from the spheroid centroid. Each point represents the median nucleus volume of a single spheroid. Lines connect paired inner and outer layer medians from the same spheroid. Black horizontal bars indicate the group median. Paired differences were assessed with a Wilcoxon signed-rank test (small: $p = 0.44$, large: $p = 0.31$).

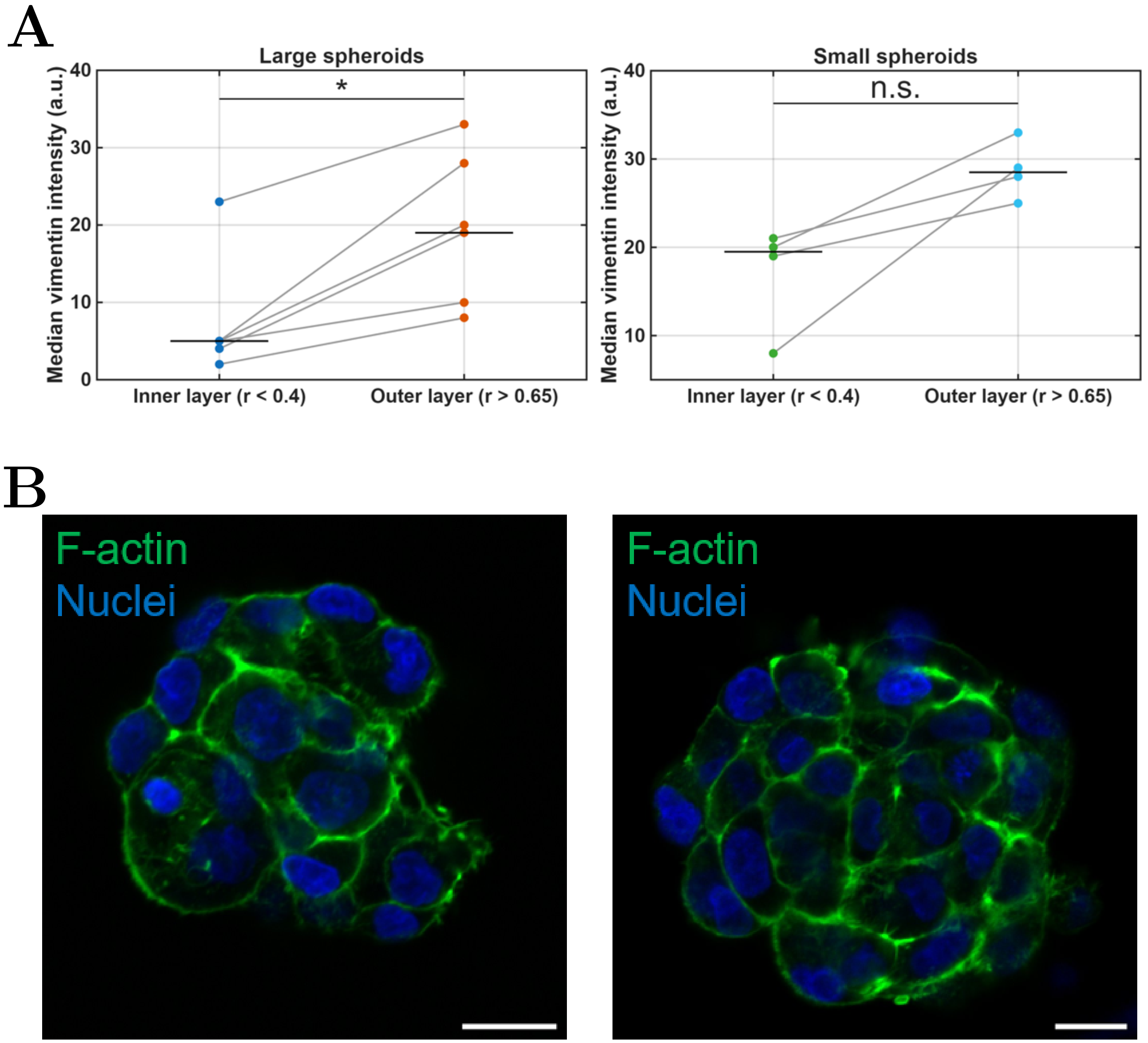


Figure S14: **(A)** Median vimentin intensity in inner layer (normalized radial distance, $r/D_0 < 0.4$) and outer layer ($r/D_0 > 0.65$) of the analyzed small and large spheroids. A significant difference in median vimentin intensity between the inner and outer layers was observed in large spheroids ($N = 6$, $p = 0.016$), whereas no significant difference was observed in small spheroids ($N = 4$, $p = 0.125$). Paired differences were assessed with a Wilcoxon signed-rank test. **(B)** Representative immunofluorescence confocal images of F-actin and nuclei from two spheroids of different sizes (left image: a small spheroid with diameter of 73 µm, and right image: a large spheroid with diameter of 105 µm), showing mid-depth optical sections. Scale bar is 20 µm.
